# Strong and localized recurrence controls dimensionality of neural activity across brain areas

**DOI:** 10.1101/2020.11.02.365072

**Authors:** David Dahmen, Stefano Recanatesi, Xiaoxuan Jia, Gabriel K. Ocker, Luke Campagnola, Stephanie Seeman, Tim Jarsky, Moritz Helias, Eric Shea-Brown

## Abstract

The brain contains an astronomical number of neurons, but it is their collective activity that underlies brain function. The number of degrees of freedom that this collective activity explores – its dimensionality – is therefore a fundamental signature of neural dynamics and computation (1–7). However, it is not known what controls this dimensionality in the biological brain – and in particular whether and how recurrent synaptic networks play a role (8–10). Through analysis of high-density Neuropixels recordings (11), we argue that areas across the mouse cortex operate in a *sensitive regime* that gives these synaptic networks a very strong role in controlling dimensionality. We show that this control is expressed across time, as cortical activity transitions among states with different dimensionalities. Moreover, we show that the control is mediated through highly tractable features of synaptic networks. We then analyze these key features via a massive synaptic physiology dataset (12). Quantifying these features in terms of cell-type specific network motifs, we find that the synaptic patterns that impact dimensionality are prevalent in both mouse and human brains. Thus local circuitry scales up systematically to help control the degrees of freedom that brain networks may explore and exploit.

## Introduction

The complexity of a neural network’s activity can be measured by its dimensionality – that is, the number of collective degrees of freedom that its neurons explore. Dimensionality is closely linked to neural computation. Signal classification, for example, benefits from network activities that increase the dimensionality of the incoming signals to be classified (2–4, 13, 14). However, compressing inputs into lower-dimensional activity patterns helps generalization to novel signals (1, 15, 16). Studies have emphasized the comparatively high (3, 6, 7) or low (17, 18) dimensionality of recordings in various experimental settings. Moreover, the dimensionality of neural dynamics can change over time (19), throughout the information processing hierarchy (20), or during learning (15, 21, 22). These findings, taken together, highlight the importance of dimensionality as a property of network activity that will vary depending on the type of computation performed in a circuit. A key question is: how can the connectivity of a network regulate the dimensionality of its activity (8, 9, 23–25)?

This question is of particular interest for cortical networks that typically exhibit quite asynchronous activity (26–28); this asynchrony at first seems to imply high dimensional dynamics with all neurons being roughly independent. However, by analyzing electrophysiological recordings from over 8000 neurons (11, 29) we show that the opposite is the case: dynamics are constrained to spaces of very low dimension relative to the number of neurons in areas across the brain. This results from the rapid accumulation of many weak but diverse pairwise correlations across the networks (cf. (30)) – as quantified by the variance of these correlations, over and above their average.

To understand the mechanistic origins of this low relative dimensionality of cortical activity, we show how the corresponding high variance of correlations results from the network’s strong recurrent connectivity. Moreover, we show that in this strongly recurrent regime, dimensionality is highly sensitive to changes in recurrent connectivity. Beyond overall synaptic strength, specific connectivity patterns, or motifs, between pairs and triplets of cells (25, 31–39) can also tune the dimensionality of neural activity. We analyze newly released synaptic physiology datasets (12, 40), quantifying connections among more than 32,000 pairs of neurons, and find that the connectivity motifs implicated by our theory are strongly present in both mouse and human brain. Moreover, connectivity strengths and motifs depend on the activity level of cell types engaged in a circuit at a given time, providing a mechanism by which dimensionality can change across cortical states in line with marked dimensionality transitions we observe in cortex.

### Highly constrained dimensionality of neural states

To quantify dimensionality in vivo, we analyzed large-scale electrophysiology recordings from the mouse visual cortex, part of the Allen Brain Observatory (11, 29). In each experimental session neural activity was recorded with six Neuropixels probes (from N = 125-520 neurons per session across the visual cortex, 319±86 on average across 26 recorded sessions) while head-fixed mice freely behaved on a treadmill with no visual stimulus (gray mean-luminance screen condition). We first observed that this activity appeared to follow a sequence of distinct states (41, 42), in which periods of roughly stationary neural activity were interrupted by abrupt changes (Figs. 1b to 1c, see Suppl. Video) This suggests that dimensionality should be quantified within and across states, rather than as a single all-encompassing number.

**Fig. 1.**
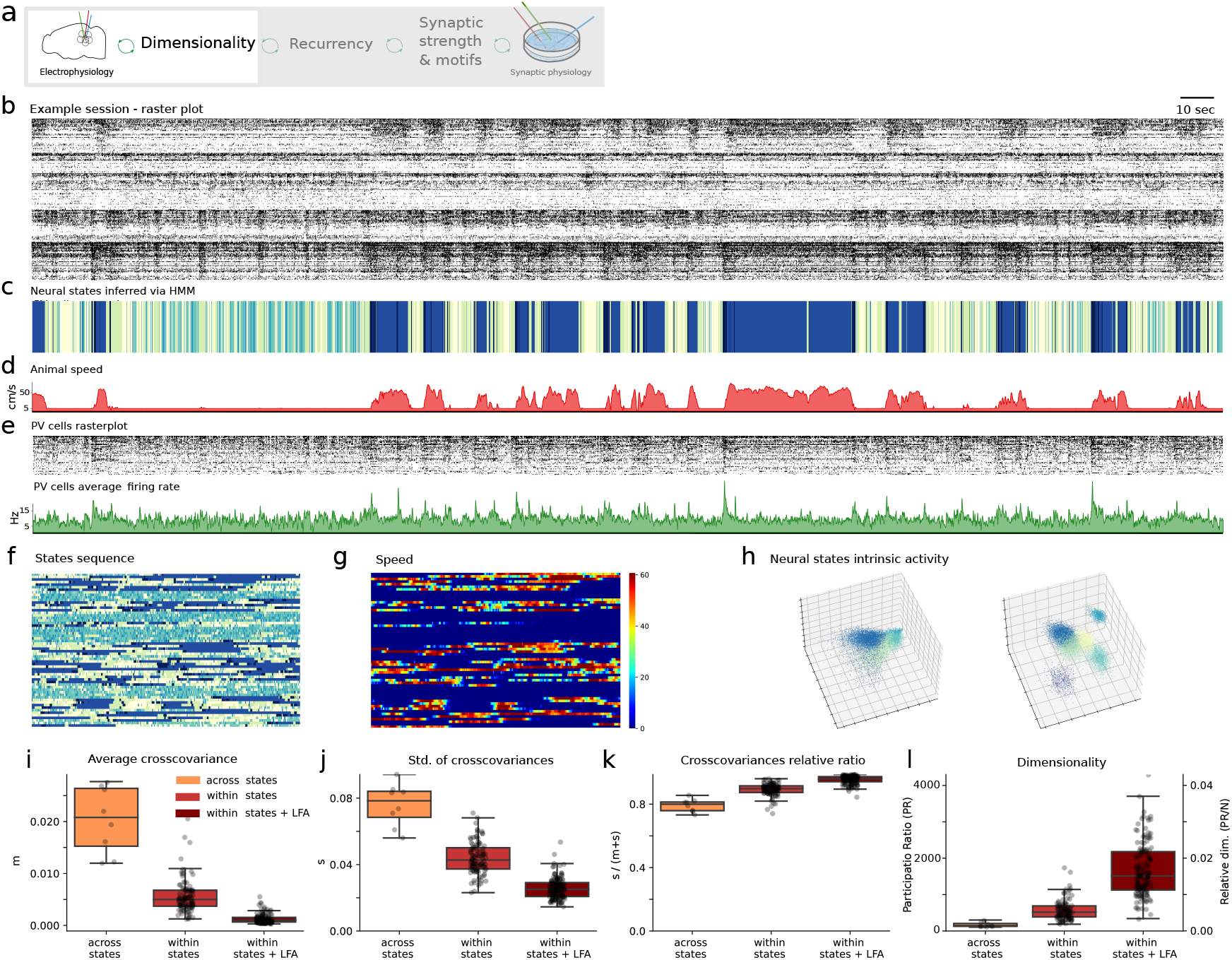
Dimensionality estimation in neural circuits. a) Figure focus: Dimensionality inferred via electrophysiology recordings. b) Raster plot example of Neuropixels recordings for one experimental session (session id=715093703). c) Sequence of states inferred by the HMM. d) Animal speed. e) Top panel: raster plot of parvalbumin (PV) cells. Bottom panel: average firing rate of parvalbumin cells. f) Neural states across the entire recording session. Each row represents a minute of activity from top to bottom, in parallel to the recording. Color represents the HMM inferred state at each given time. g) Same as f where color represents the animal speed. h) Neural activity of example session for each neural state before (left) and after (right) performing LFA (Latent Factor Analysis) on the activity of each neural state. The three axes represent directions of the top three PCs after performing PCA (Principal Components Analysis); same color code as in c and f. i) Normalized cross-covariances 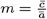 (see explanation of Eq. (1)) averaged across neurons and across the entire experimental session (orange), averaged within each neural state (red) and after performing LFA for each state (bordeaux). j) Same as i but for the normalized standard deviation of cross-covariances 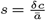. k) Same as i for the relative ratio (*s/*(*s* + *m*)). l) Same as i for the dimensionality measured by the participation ratio (PR), Eq. (1). Left scale (y-axis) extrapolated dimensionality to 10^5^ neurons, right scale (y-axis) in percentage normalized to the number of neurons.

We identified states using a Hidden Markov Model (HMM). Here, transitions among states are assumed to occur discretely via a Markov chain; within each state (colored intervals in Fig. 1c and Fig. 1f), neurons fire spikes as Poisson processes, with firing rates particular to that state. To restrict to clearly identified states, we retained only time bins for which the likelihood of being in the identified state (HMM model’s posterior probability) was at least 80% (time bins where no state satisfied this criterion were left unassigned). In each session the number of states was set via an unsupervised cross-validation procedure (7.4 ± 1.7 across 26 sessions, which ranged from 5 to 11 patterns, see Methods and Fig. S1). Neural states appeared separated in the space of population responses (Fig. 1h), and were influenced by animal behavior (e.g. running, Fig. 1d).

We computed the widely used participation ratio *D*_PR_ (17, 19) (Fig. S2) to measure dimensionality. *D*_PR_ is defined via the percentages of explained variance of activity along the principal components, given by the eigenvalues of the covariance matrix of neural activity. Importantly, *D*_PR_ can also be expressed exactly in terms of the statistics of entries in this covariance matrix *C*, which can be tractably measured and modeled (Fig. S2b). Specifically, where *N* is the number of neurons analyzed (see Methods and (2, 19)):

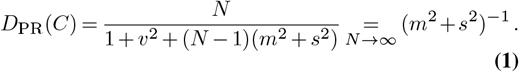

Here, *v, m* and *s* are respectively the ratios between the standard deviation of variances (*δa*), the average of cross-covariances 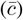 and the standard deviation of cross-covariances (*δc*), to the average of auto-covariances 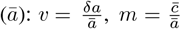 and 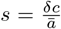. Here, *ā* may be thought of as a firing rate normalization. Eq. (1) shows that, for large populations *N*, the factors *m* and *s* are the key factors modulating dimensionality. While each appears equally in the formula, the two terms may have largely different origins. For example, a high *m* can be driven by a relatively small number of “low-rank” behavioral components (7, 43) while a high *s*, as we will show, may result from strong recurrent connections among neurons.

Thus, we compute *m* and *s* in the neural recordings, and ask whether either dominates. We first compute values across the entire period of activity (i.e., across states), and then separately within each neural state (Fig. 1i, Fig. 1j). As expected due to state transitions, *m* was significantly greater across states than within them. The same was true for *s*, although to a much lesser extent. Overall, *s* dominates *m* (Fig. 1k), with the ratio *s/*(*s* + *m*) exceeding 90% within states. This is consistent with previous findings for cortical activity data (27, 28) and models of cortical networks (23, 26, 28, 44, 45). We verified that this trend becomes even stronger when we remove remaining low-rank components of the covariance via a cross-validated Latent Factor Analysis (LFA) for each neural state (Fig. S3, Fig. S4).

Finally, we combined the contributions of *m* and *s* (together with *v*) to compute the dimensionality *D*_PR_(*C*) (Eq. (1)). Exploiting the explicit dependence of dimensionality on the number of neurons *N*, we developed a systematic extrapolation to larger sample sizes *N* (see Fig. S2f, Fig. S2i, Methods). The results revealed that dimensionality had values of only ∼ 100 − 1000 dimensions even when *N* = 10^5^ – a typical reference value for the number of neurons in a local cortical network (cf. Fig. 1l). In principle, the dimensionality for a set of uncorrelated neurons can be as high as the number of neurons *N*, with each exploring a separate degree of freedom; specifying dimensionality relative to this scale shows that *D*_PR_(*C*)*/N* ⪅ 1%. This is a strikingly small fraction, and indicates highly constrained neural activity patterns. These values were consistent when computed across both cortical layers and areas (Fig. 2e) as well as with different measures of dimensionality (Fig. S7) and bin sizes (Fig. S8).

**Fig. 2.**
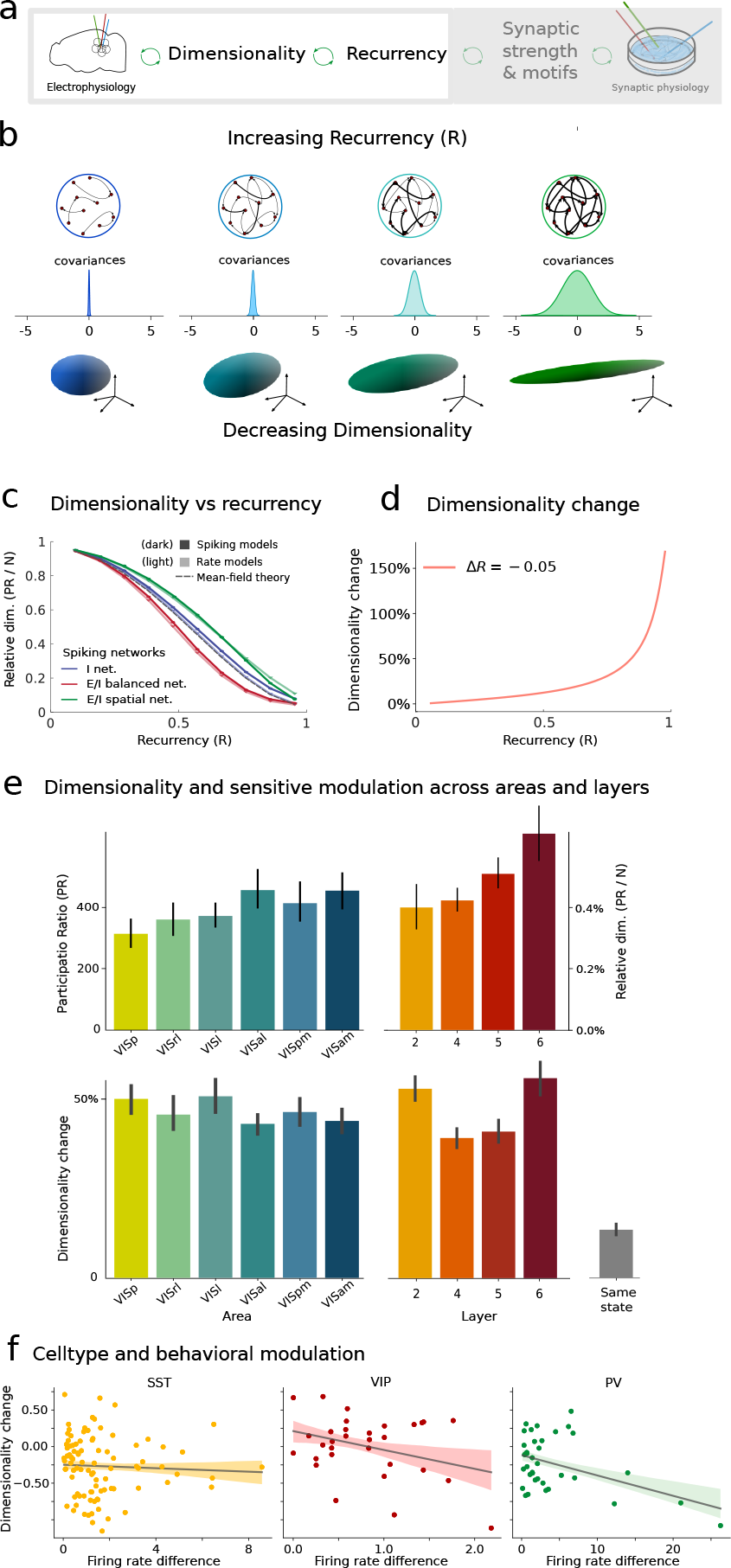
Dimensionality and recurrency *R* across the visual cortical circuit. a) Figure focus: Dimensionality is linked to recurrency. b) Three-way connection between recurrency, width of covariance distribution and dimensionality of neural activity. c) Normalized dimensionality *D*_PR_*/N* as a function of the recurrency *R*. Theoretical predictions for the dimensionality in homogeneous inhibitory networks (gray) are accurate for simulations of rate models (light colors) and spiking models (dark colors) across various network topologies (blue: homogeneous single population inhibitory networks, red: homogeneous two-population excitatory-inhibitory networks, green: spatially organized single-population inhibitory networks). d) Dimensionality change (relative modulation of dimensionality) as a function of recurrency, (Δ *D*_PR_). e) Dimensionality extrapolated to network size of 10^5^ neurons (top) and dimensionality change (relative modulation of dimensionality) between neural states (bottom) across visual cortical areas and cortical layers. Control: gray bar (rightmost plot), dimensionality change computed between multiple instances of the same neural state. f) Signed dimensionality change (signed 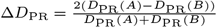) versus average firing rate difference of neurons of specific cell type (SST, VIP and PV) between pairs of neural states.

### Strong recurrence as a mechanism for low dimensionality

What is the mechanistic origin of this strongly constrained dimensionality of neural activity? We propose that it lies in the underlying recurrent network connectivity, (Fig. 2a), and how this sets the main quantity *s* determining dimensionality. Previous studies (23, 45) showed that *s* is determined by the level of network recurrency for a given neural state: as network recurrency becomes stronger, longer and longer synaptic paths impact the co-variation in activity between each pair of neurons. These longer paths are highly variable from one neuron pair to the next, and this variability drives a wide range in the cross-covariances across neural pairs, precisely the quantity that *s* measures (Fig. 2b).

This intuition, linking co-variation in neural activity to underlying synaptic paths, can be formalized by means of quantifying recurrency through a single number *R*, the spectral radius of connectivity eigenvalues. *R* summarizes the statistics of network connections and the overall strength of recurrent coupling (see Methods). Via a three-way link between low-dimensional neural activity (Fig. 2b bottom), large variance of correlations (Fig. 2b center) and strong recurrent connections (Fig. 2b top), there is a direct relationship between dimensionality and recurrency *R*:

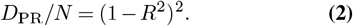

(see Suppl. Mat. S4.8 and S4.9 for a formal derivation based on (23) and (25) for an alternative derivation). This relationship is robust, as shown by our validation in nonlinear spiking networks, and holds for networks with a wide range of topologies (Fig. 2c).

In turn, the relationship between dimensionality and recurrency suggests that the low relative dimensionality of cortical activity (⪅ 1%, Fig. 1l) can be explained if visual cortical networks operate in a strongly recurrent regime (46), where *R* is close to the critical value *R* = 1. This conclusion of strong recurrency is robust under assumptions about inputs to networks (Fig. S4, Fig. S6a). Specifically, in models including inputs with relatively low-rank components shared across many neurons, we still find that strong recurrency is required to quantitatively reproduce the activity statistics observed in neural recordings (Fig. S5).

### The sensitive regime for modulation of dimensionality

In the same strongly coupled regime, the relative change in dimensionality with respect to a change Δ*R* in recurrency *R* is highest. Fig. 2d illustrates this via the quantity 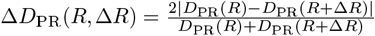. Consequently, when *R* increases, neural networks acquire a sensitive control of dimensionality: the capacity to sharply regulate dimensionality via underlying connectivity features that impact the recurrency *R*. As above, we verified that this strong sensitivity remains in the presence of low-rank network inputs consistent with the neural recordings (Fig. S6b).

Our proposal that cortex operates in a regime where dimensionality is under sensitive control suggests that different cortical states should produce notably different dimensionalities. To test this, we measured the dimensionality change between any two states *A* and *B* in the same session, 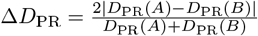. Values close to or above 50 percent for Δ*D*_PR_, as found in our analysis (Fig. 2e), imply that dimensionality changes across the two states are substantial, on average roughly half or more as large as the dimensionalities themselves.

What factors could contribute to the differences of dimensionality across states? States are defined by different neuronal firing rate patterns, raising the possibility that cell types are differentially engaged in each. Thus, we first checked whether dimensionality across states was increased when neurons of vasoactive intestinal peptide-expressing (VIP), somatostatin (SST), and parvalbumin (PV) cell types significantly increased their firing rate within a given state. Neurons were identified based on their responses to optogenetic stimulation (see Methods). We found a trend in dimensionality for PV cells, but to a lesser or no extent for VIP and SST types, respectively (Fig. 2f). These varied effects motivate the need to elucidate the mechanistic origin of different levels of recurrency, to which we turn next.

### Local tuning of the global recurrency *R*

We asked *how* neural networks can regulate their overall recurrency *R* and hence their dimensionality (Fig. 3a). While many previous studies established how global features of recurrent connectivity affect *R* (23, 47, 48), here we focus on the impact of local features of connectivity: connection probabilities and strengths and second order connectivity motifs. These motifs are statistics of the neural connectivity *W* that involve *pairs* of connections (see Methods), and together with overall connection probabilities form the fundamental local building blocks of networks. Second order motifs appear in four types (Fig. 3b): reciprocal, divergent, convergent, and chain motifs, together with the variance (strength) of neural connections already present in purely random models (47). These motifs have been shown to play important roles determining neuron-to-neuron correlations and allied circuit dynamics (31–34, 37, 49–53) and emerge from learning rules consistent with biological STDP mechanisms (54, 55).

**Fig. 3.**
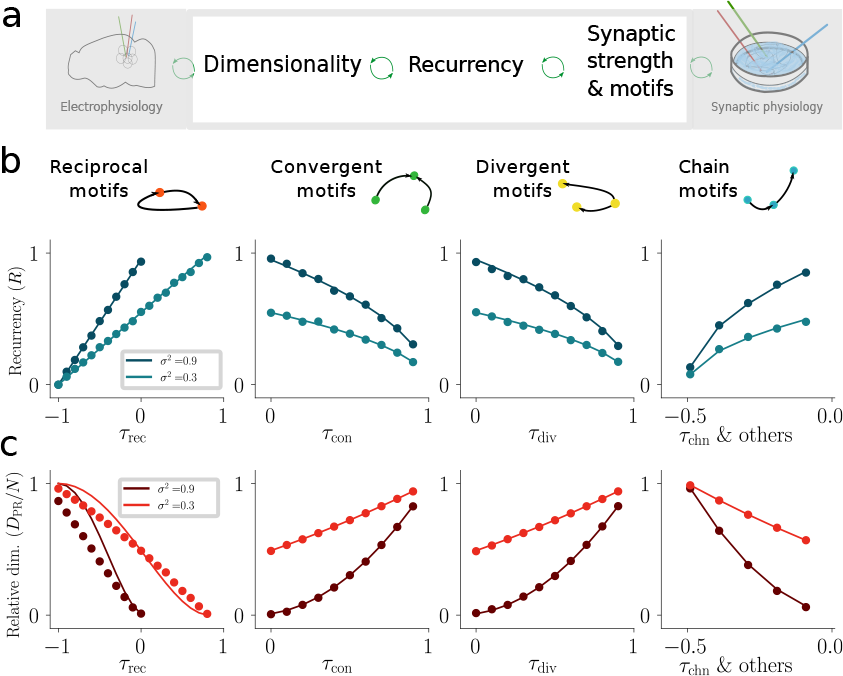
Theory for recurrency and dimensionality in networks with second-order motifs. a) Figure focus: Modulation of recurrency and dimensionality by local circuit motifs. b) Theoretical dependence of recurrency on abundances of reciprocal, convergent, divergent and chain motifs. c) Theoretical dependence of dimensionality on motif abundances. Solid lines: theory. Markers: simulations. The abundance of chain motifs *τ*_*chn*_ is constrained by theoretical limits, as reflected in the range of values of *τ*_*chn*_ (See Suppl. Mat. S4.11).

We developed a comprehensive theory that takes full account of all second order motifs in networks of excitatory and inhibitory neurons, generalizing allied results developed via distinct theoretical tools (25, 52). Our analysis yields a novel compact analytical quantity that shows how recurrency is modulated by local structure: *R* = *σ* · *R*_motifs_.

The first term, *σ*, stems from the strength and number of connections and has been studied before (56) (see Suppl. Mat. S4.7). The second:

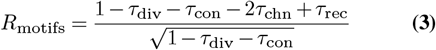

compactly describes the influence of connectivity motifs, in a form that is novel here. In this equation *τ*_rec_, *τ*_chn_, *τ*_div_, *τ*_con_ respectively capture the abundance of reciprocal, chain, divergent, and convergent motifs (cf. Methods and Suppl. Mat. S4.8). This formula describes how the recurrency *R* is affected by increasing or decreasing the prevalence of specific motifs (Fig. 3b) and thus links the modulation of auto- and cross-covariances and the dimensionality of neural responses across the global network to the statistics of local circuit connectivity, as shown in Fig. 3c. While Eq. (3) is exact for networks with homogeneous connection statistics, we show that it generalizes to models with distinct connection statistics for excitatory and multiple inhibitory neuron types. Here, *σ* and *τ* combine the corresponding statistics of the excitatory and inhibitory populations (cf. Fig. S9 and Suppl. Mat. S4.9). This direct link between quantifiable, local connectivity statistics and the global network property *R* opens the door to novel functional analyses of very large-scale synaptic physiology datasets in both mouse and human, as we describe next.

### Cortical circuits in mouse and human employ local synaptic motifs to modulate their recurrent coupling

We analyzed newly released synaptic physiology datasets from both mouse and human cortex (12, 40) to assess the involvement of synaptic connectivity properties in modulating network recurrency and to probe their possible role in driving the changes in dimensionality across states summarized in Figs. 2e to 2f. This dataset is based on simultaneous in-vitro recordings of 3-to-8 cell groups (cf. Methods) and consisted of 2, 767 identified synapses from mouse primary visual cortex (out of more than 32, 000 potential connections that were tested) and 363 synapses from human cortex.

Recall that the recurrency *R* as defined above has an overall scaling term, *σ*, and a motif contribution term given by Eq. (3). We began by computing *σ*, which is determined by the probabilities and strengths of individual connections within the mouse V1 network, and how these differ across layers and cell types (see Suppl. Mat. S4.7). First, we observed that neurons within and across the excitatory and inhibitory populations of layer 2 are on average more connected than those of layer 5 (Fig. 4b). This predicts a higher *σ* and hence *R* in layer 2 (Fig. 4d), consistent with the lower dimensionality found there (Fig. 2e right). Second, as in (12), we noted that PV cells are more connected to pyramidal cells, compared with SST and VIP cells (Fig. 4c). This predicts that changes in PV activity will have a greater effect on recurrency and hence dimensionality (Fig. 4e, consistent with Fig. 2f).

**Fig. 4.**
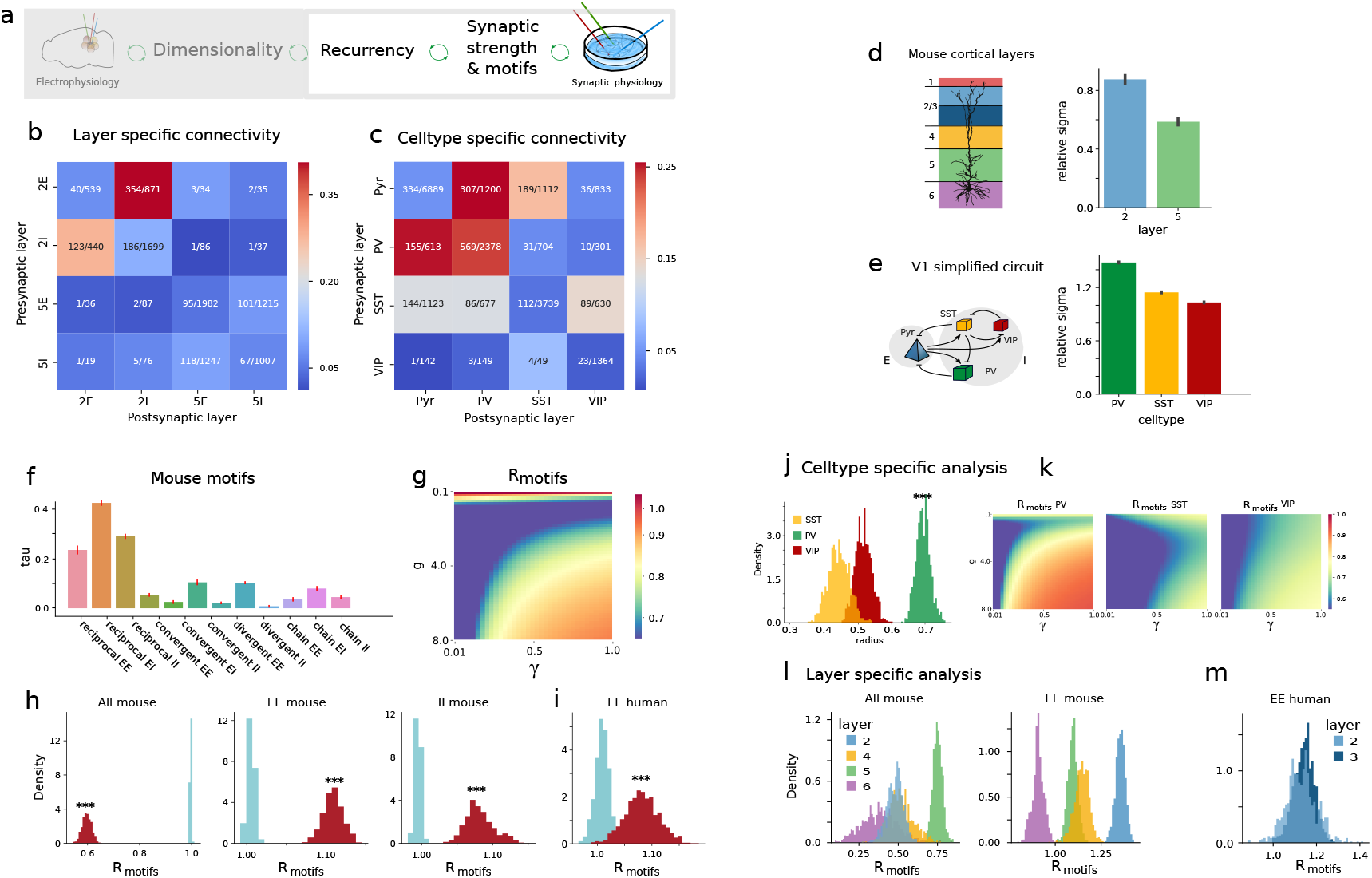
Motif analysis in synaptic physiology datasets. a) Figure focus: network recurrency inferred via synaptic physiology datasets. b) Statistics of mouse connectivity across layers for excitatory and inhibitory synapses (synapses identified / synapses probed). c) Statistics of mouse connectivity across cell types. d) Relative strength of connectivity for layer 2 and 5 in mouse V1 (*σ*_LX_ */ σ*_L2+L5_ for *X* ∈ {2, 5}, see Suppl. Mat. S4.7). e) Relative strength of connectivity for mouse V1 circuit with multiple cell types (*σ*_X_ */ σ*_PV+SST+VIP_ for *X* ∈ {*PV, SST, V IP*}). f) Motif abundances in mouse V1. g) Inferred *R*_motifs_ as a function of relative strength *g* of inhibitory and excitatory synapses and ratio *γ* of inhibitory to excitatory population size. Unless stated otherwise, the reference values used in the analyses are *g* = 4, *γ* = 0.5. h) Estimation of *R*_motifs_ from mouse data, red distribution (500 bootstraps based on random subsets of 80% of sessions). Blue distribution: estimation of *R*_motifs_ on 500 ensembles of randomly connected neurons, see Methods. Left panel shows *R*_motifs_ computed using all synapses, both excitatory and inhibitory. Middle and right plot consider either only excitatory, or only inhibitory synapses (EE and II respectively). i) Same as h for EE motifs in human dataset. j) Inferred *R*_motifs_ for strengthened SST (left) and PV (right) populations (see Methods). k) Same as g where PV (left) or SST (right) synapses have been strengthened to test the impact of these connections on *R*_motifs_. l) Inferred *R*_motifs_ for layer specific neural populations. The computation includes synapses of a given layer that are both excitatory and inhibitory (left) or only excitatory (right). m) Layer-wise estimation of *R*_motifs_ for EE-only motifs in human.

Next, we analyzed the probability of occurrence of connection motifs, hence estimating the remaining contribution to recurrency: *R*_motifs_ (cf. Methods and Suppl. Mat. S4.8). Beginning with the mouse data, we calculated the statistics of individual motifs, separating those involving excitatory (E) and inhibitory (I) synapses (EE, EI, II, Fig. 4f), and found many motifs to be significantly present. We then combined these to compute *R*_motifs_. This requires two parameters: one regulating the overall ratio of inhibitory to excitatory neurons (*γ*), and another the relative strength of the inhibitory synapses (*g*) (cf. Suppl. Mat. S4.9). We found that *R*_motifs_ *<* 1 across almost all choices of these parameters (Fig. 4g), so that the overall role of the observed motifs is to decrease recurrency and hence increase dimensionality. Different motifs, distinctly involving E and I cell types, combine to produce this value of *R*_motifs_. To study how this occurs, we separated the contribution to *R*_motifs_ from motifs within the excitatory population (EE type only) by assuming that other motifs occur at chance level. Interestingly, the EE motifs operating alone produced the opposite trend, increasing the radius 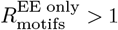 (Fig. 4h center, one-sided t-test p-value*<* 10^−20^). The same was true for motifs within the inhibitory population 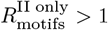 (Fig. 4h right, one-sided t-test p-value*<* 10^−20^), and for motifs within the excitatory population in human cortical circuits (Fig. 4i, one-sided t-test p-value*<* 10^−20^). We further confirmed that this effect is also predicted for previously published data on excitatory connections in rat visual cortex (35) (cf. Methods). The increased recurrent coupling strengths within both the excitatory and inhibitory populations underscore the prominent role of EI motifs, specifically reciprocal EI motifs, because these counteract the influence of EE and II motifs and lead to the overall decrease in recurrency *R*_motifs_ *<* 1 (Fig. 4h, Fig. S9).

The distinct roles of E and I motifs in regulating *R*_motifs_ point to additional ways that recurrency, and hence dimensionality, may be controlled dynamically in neural circuits. As noted above, one pathway for this control is via a differential activation of cell types (Fig. 4e), engaging different overall connection strengths *σ* in the network. Here, we explore how motifs may play an additional role in this process.

First, we separately quantified cell-type specific motifs in the synaptic physiology dataset and found strong differences across the analyzed cell types (Fig. S10). To test whether PV, SST and VIP activity modulation could alter recurrency by modulating the motifs engaged in cortical circuits, we compared the values of the motif contribution to recurrency, *R*_motifs_, for three distinct scenarios: when either the PV, the SST or the VIP is the dominant inhibitory population. We found that a circuit with a dominant PV inhibitory population had a substantially higher level of *R*_motifs_ compared to SST and VIP (Fig. 4j). This was true across all choices of parameters (Fig. 4k). This result shows an additional mechanism by which an increased activity of PV interneurons may reduce the overall dimensionality, as found in our analysis (Fig. 2f), by differentially engaging circuit motifs. In sum, our analysis shows how cell-type specific control of interneural populations can lead to a global modulation of dimensionality via local circuit mechanisms.

Next, we asked whether cross-layer differences in connectivity motifs could likewise affect the recurrency, and hence dimensionality and its modulation across states. We found that the corresponding motif contribution *R*_motifs_ for excitatory synapses was significantly stronger for layer 2 and 5 than for layer 6 (Fig. 4l). However, when combined with II and EI effects, the overall difference in *R*_motifs_ across layers 2 and 5 reversed (Fig. 4l left) for our default parameters. We conclude that we have evidence that differences in connection strength *σ* do contribute to *R* varying across layers, but in opposite effect to the differences in synaptic motifs. This is in contrast to the case for cell type modulation, for which differences in both connection strength and synaptic motifs appear to play a consistent role in changes in *R* and hence changes in dimensionality. Finally, we performed a similar analysis on the human synaptic dataset. While the more limited data allowed for fewer comparisons, this did confirm that the values of *R*_motifs_ for excitatory connections in human cortex layers 2 and 3 are broadly consistent with those observed in layer 2/3 of the mouse cortex (Fig. 4m).

## Summary and discussion

We showed that neural networks across the mouse cortex operate in a strongly recurrent regime, in which the dimensionality of their activity is strongly constrained. A feature of circuits in this regime is the ability to sensitively modulate the relative dimensionality of their activity patterns via their recurrency *R*, a unifying measure of a network’s overall recurrent coupling strength. We note that related concepts of activity sensitivity have been studied in other close-to-critical dynamical regimes, such as the reverberating regime controlled by E-I balance rather than recurrency and hypothesized to have important consequences for computation (57).

Our theoretical work links observations on dimensionality to predictions for network recurrency *R*: a higher dimensionality suggests a lower recurrency and vice-versa. Moreover, we showed that the critical circuit features that determine a circuit’s recurrency *R* are not just its overall connection strength, but also a tractable set of local synaptic motifs. We use theoretical tools to quantify the effect of these motifs via a compact index *R*_motifs_. This provides a concrete target quantity that can, as we show, be readily measured from emerging, large-scale synaptic connectivity datasets and used to test predictions about the role of synaptic structure in controlling dimensionality. We show how the engagement of different inhibitory cell types (58, 59) form one avenue for controlling dimensionality via local connectivity statistics. This is, of course, the tip of the iceberg in terms of how cortical circuitry is modulated across time and state, and allied effects are likely prominent. As we reviewed above, high dimensional activity can retain stimulus details, while lower dimensional activity can promote robust and general downstream decoding; taken together, this points to new functional roles for modulatory and adaptive mechanisms known to take effect across time and across brain states.

In sum, our theory and brain-wide experimental analyses converge to provide new evidence for an intriguing concept (52, 60, 61): that the connectivity of cortical brain networks exerts control over their activity in a highly tractable manner, via the building blocks of their local circuitry.

## Acknowledgments

We thank Krešimir Josić, Brent Doiron, Luca Mazzucato, Stefan Mihalas, Nicholas Steinmetz, Michael Okun, Byron Yu, Daniel Denman, Yu Hu, Severine Durand and Leenoy Meshulam for helpful discussions. S.R. was supported by a Swartz Fellowship in Theoretical Neuroscience at the University of Washington, and by NIH BRAIN Grant R01EB026908, and E.S.B. by NIH R01EB026908, NIH BRAIN 1RF1DA055669 and NSF Grants DMS 1514743 and NCS-FO 2024364. D.D. and M.H. were supported by the HGF young investigator’s group VH-NG-1028, the European Union’s Horizon2020 research and innovation program under Grant agreements No. 785907 (Human Brain Project SGA2) and No.945539 (Human Brain Project SGA3), and funded under the Excellence Strategy of the Federal Government and the Länder (G:(DE-82)EXS-PF-JARA-SDS005). We thank the Allen Institute for Brain Science founder, Paul G. Allen, for his vision, encouragement, and support.

## Data availability

The data that support the findings of this study are available from the corresponding author upon reasonable request.

## Code availability

The code for the numerical simulations and data analyses are immediately available from the corresponding author upon reasonable request, and will be released in a public repository before final publication of the manuscript.

