## Supplementary material for "Strong and localized recurrence controls dimensionality of neural activity across brain areas": Methods and Supplementary Materials

Supplementary Figures, Methods and Theoretical Supplementary Material

David Dahmen, Stefano Recanatesi, Xiaoxuan Jia, Gabriel K. Ocker, Luke Campagnola, Stephanie Seeman, Tim Jarsky, Moritz Helias, Eric Shea-Brown

**5 Bibliography** **39**

### 1 Structure of Supplementary Figures

We first provide a concise summary of the structure and motivation of the supplementary figures in order to facilitate access to these figures, as well as the more detailed supplemental analyses that follow.

Fig. S1 provides further details on the Hidden Markov Model (HMM) analysis used to infer various neural states from the spontaneous activity of the mouse visual cortex. This HMM analysis is the basis for the results presented in Fig. 1 and Fig. 2 of the main text. For more details, see Sec. 3.1.2.

Fig. S2 introduces the concept of dimensionality of neural activity, its relationship to the statistics (means and standard deviations) of activity covariances, and a technique for extrapolating to larger ensembles of neurons. We demonstrate on model data that this extrapolation method produces unbiased estimates of the dimensionality of the entire network from the activity of a subset of neurons. This subsampling reflects the experimental situation, in which we record the activity of only hundreds of neurons embedded within much larger networks. The results of the dimensionality extrapolation based on experimentally measured statistics are the basis for the dimensionality analyses in Fig. 1 and Fig. 2 of the main text. For more details, see Sec. 4.1 and Sec. 4.2.

Fig. S3 illustrates the cross-validated Latent Factor Analysis (LFA) method applied to the activity within various neural states, in order to extract low-rank components that could have originated from external inputs. This analysis validates our results on strongly constrained dimensionality of the experimentally recorded neural activity against the potential influence of external inputs. In Fig. 1 of the main text, the resulting statistics of covariances and dimensionality of activity are compared with and without removal of low-rank components. In addition, the figure illustrates how we eliminate the bias in the covariance statistics due to a finite number of samples (time bins) and obtain an unbiased estimate of the cross-covariance standard deviation (for details, see Sec. 4.3), as is needed to obtain an unbiased estimate of the dimensionality.

Fig. S4 applies the cross-validated LFA procedure to model data generated by simulating a network model. This allows us to verify whether the cross-validated LFA technique recovers the known ground truth. It shows that LFA faithfully removes externally driven low-rank components of activity, but also tends to remove strong intrinsically generated activity components that arise from local reverberation of fluctuations via strong recurrent connections. Consequently, the recurrency inferred from simulated activity is typically underestimated, resulting in an overestimation of the the inferred dimensionality of intrinsic activity, especially in the strongly recurrent regime. Therefore, we show that the LFA procedures yield a conservative estimate of how low the dimensionality of intrinsic activity is. The dimensionality values shown in Fig. 1l (within states + LFA) can therefore be interpreted as upper bounds to the true values. For more details, see Sec. 4.5.

Fig. S5 demonstrates that the network model used in our theoretical study, when fit to the experimentally measured mean and standard deviation of auto- and cross-covariances, can accurately describe the experimental activity data in terms of the covariance eigenvalue spectra and dimensionality. For networks of reasonable size, the fit model must also operate in the strongly recurrent regime in order to adequately explain the data, even in the extreme case where all low-rank activity components are attributed to external inputs. Overall, the analysis indicates that a linear network model operating in the strongly recurrent regime is a minimal but adequate model for capturing the experimentally observed covariance statistics and dimensionality. For more details, see Sec. 4.6.

Fig. S6 shows the dependence of dimensionality on network recurrency in the presence of experimentally realistic external input. Network and input parameters are first inferred from experimental activity data from some example HMM states. Next the recurrency $R$ of the fit models is modulated as in Fig. 2c,d of the main text, to assess its role in determining dimensionality. We find that while external input and distributed firing rates lead to an overall reduction in dimensionality, network recurrency further constrains the activity in the strongly recurrent regime and leads to a high sensitivity with respect to changes in $R$. This analysis thereby serves as a control to confirm that the relation between low/sensitive dimensionality and strong recurrency shown in Fig. 2c,d in the main text is also valid in the presence of external experimentally realistic inputs. For more details, see Sec. 4.6.

Fig. S7 compares our results using the participation ratio measure of dimensionality, with results based on other measures of dimensionality. The participation ratio measure is shown to correlate well with a measure based on the number of components required to explain 70% of the variance of neural activity. Additionally, if not extrapolated according to our method (Eq. S1, see also Sec. 4.2), the participation ratio as well as these other measures, shows a strong dependence on the number of recorded neurons. However, this dependence of the participation ration on the number of recorded neurons does not appear substantial once the dimensionality has been extrapolated according to our method, provided that enough data is recorded to begin with. We conclude that the extrapolated participation ratio is therefore a more appropriate method for comparing distinct sessions, regions, and layers, as shown in Fig. 1 and Fig. 2 of the main text.

Fig. S8 shows the dependence of dimensionality (extrapolated participation ratio) on the sizes of time bins in which spikes are counted, across different visual areas and cortical layers. The results are generally consistent across bin sizes of 50ms, 100ms and 200ms.

Fig. S9 shows a validation of our theoretical predictions (see Sec. 4.9) for the recurrency (spectral radius) of excitatory-inhibitory networks with population specific abundances of reciprocal, convergent, divergent and chain motifs, against numerical analyses of connectivity matrices generated with the algorithm described in Sec. 4.11. The figure is analogous to Fig. 3b of the main text, but for the case of two populations.

Fig. S10 shows that the abundance of reciprocal, convergent, divergent and chain motifs of pyramidal cells with different inhibitory cell types varies strongly in the synaptic physiology dataset, indicating that differential activation of inhibitory cell types can have differential impacts on the overall recurrency of the underlying network, as shown in Fig. 4j,k.

#### 2 Supplementary Figures

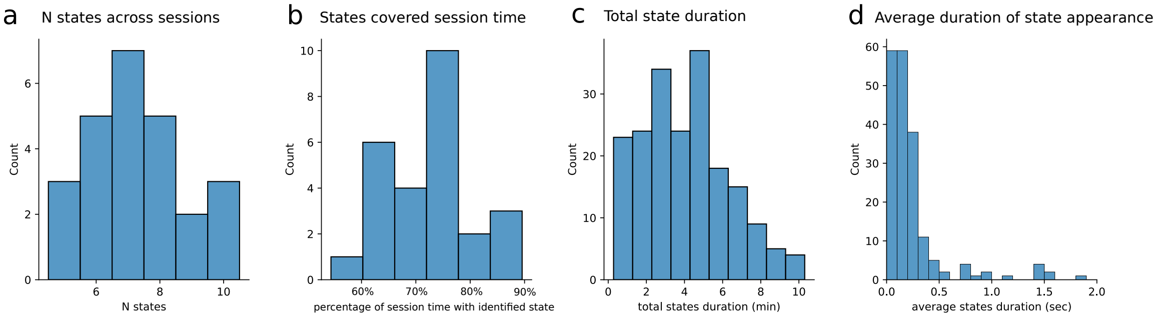

Figure S1: **Hidden Markov Model (HMM) analysis of spontaneous activity in mouse visual cortex.** The HMM algorithm (see Methods) extracts, in an unsupervised way, periods where the neural activity is well captured by the same latent state. It returns a parsing of 30 minutes of spontaneous periods of activity into multiple underlying states. In each state the hypothesis of the algorithm is that the average neural activity is steady. a) Histogram of number of detected states across all recording sessions. b) Histogram across all sessions of the fraction of session time (in percentage) covered by HMM states detected above the confidence threshold. c) Histogram of total durations, across all appearances of an individual state within a session, of states across all recording sessions. d) Histogram of average duration of individual occurrences of HMM states across all states and sessions.

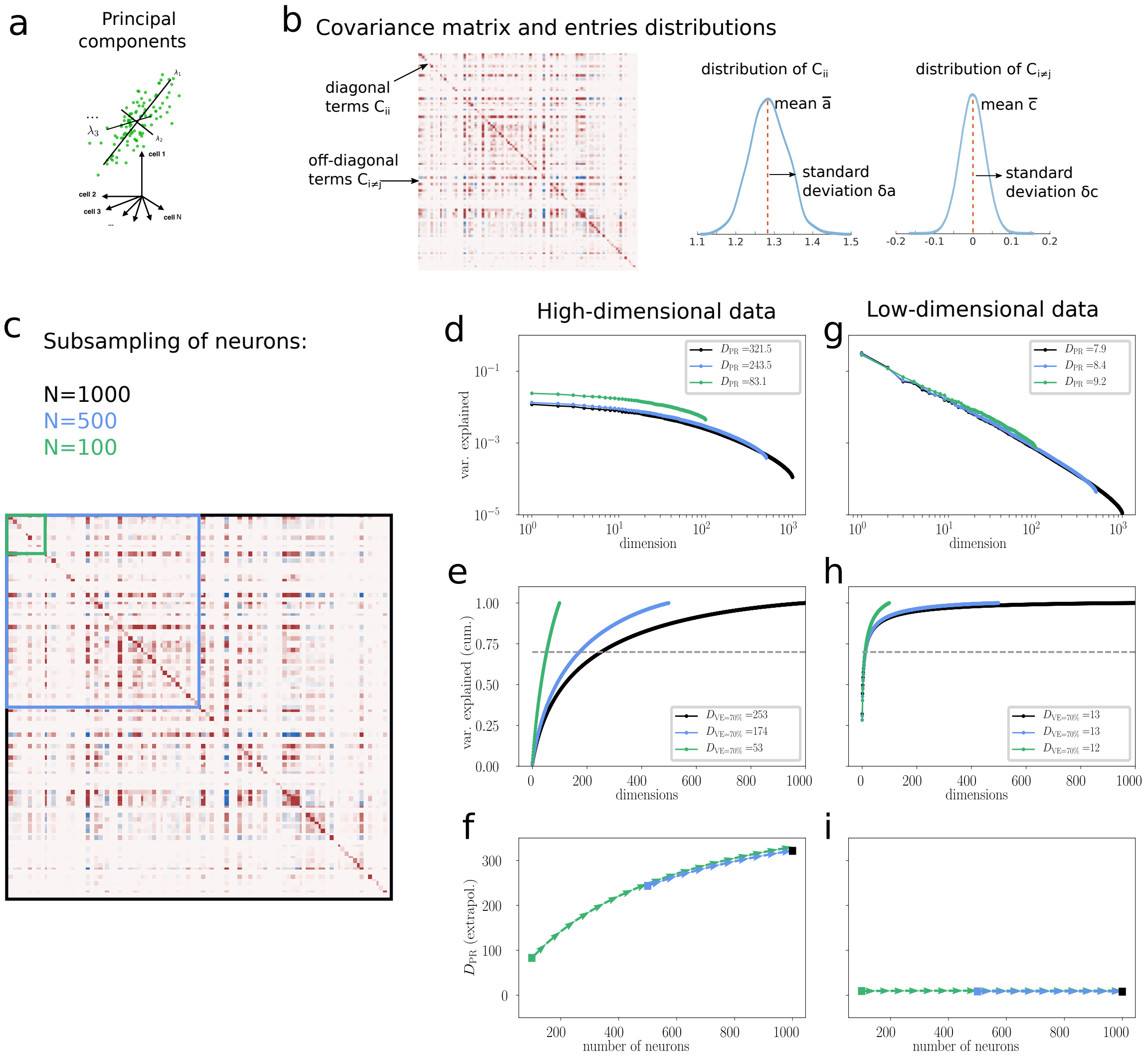

Figure S2: **Statistics of covariances and dimensionality estimation.** a) Example of data distribution where each green dot corresponds to a singlevector of activity. In our experimental analyses, such a vector contains the spike counts of all neurons in a window of 100ms duration. The geometrical interpretation for such a point cloud is that the axes of major variability are determined by the eigenvectors of the covariance matrix. Eigenvalues, when normalized to their sum, determine the amount of variance of the data in the direction identified by the eigenvector. b) Example of spike-count covariance matrix and statistics. The two distributions on the right capture respectively the statistics of diagonal (auto-covariances) and off-diagonal (cross-covariances) entries of the covariance matrix. Of specific importance for our study are the standard deviation of cross-covariances $\delta c$ and the mean of auto-covariances $\bar{a}$ as their ratio, here termed $s$, is in one-to-one correspondence with the recurrency (spectral radius) $R$. c) Illustration of the subsampling analysis performed in panels d-i, where only subset of neurons is taken into account for the analysis (black: $N = 1000$ neurons, blue: $N = 500$ neurons, green: $N = 100$ neurons). d-f) High-dimensional activity data from a model simulation. d) Variance explained of individual activity components as inferred via principle component analysis (PCA). The shape of this spectrum varies with the number of neurons that are included in the analysis. Consequently, the value of participation ratio $D_{\mathrm{PR}}$, which is based on this spectrum, varies strongly. e) Cumulative variance explained as a function of the number of principal components. The same level of variance explained (here 70%) is obtained from largely different numbers of components for the different subsamplings. f) Although the participation ratio $D_{\mathrm{PR}}$ yields different values for the different subsamplings, Eq. S1 allows the correct inference of the dimensionality of larger datasets, in particular the full network of $N = 1000$ (black marker), by means of extrapolation. g-i) Same as d-f, but for low-dimensional activity data. Since the data is much lower dimensional than all the subsamplings, the inferred dimensionality values via $D_{\mathrm{PR}}$ and cumulative variance explained curves are more consistent. The extrapolation procedure of $D_{\mathrm{PR}}$ can still be applied but does not have any sizable effect.

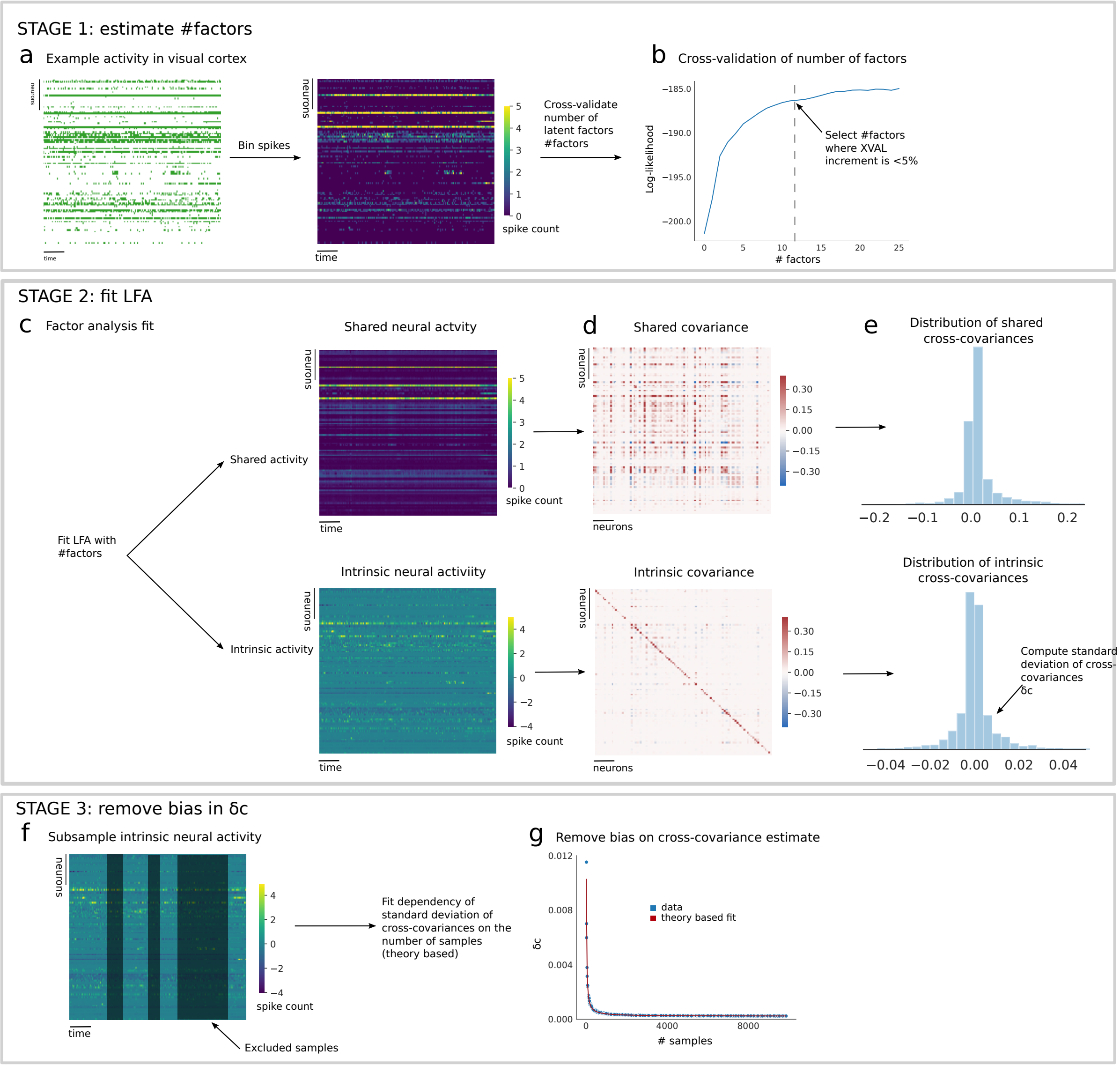

Figure S3: **Method for removing shared low-rank activity components to infer intrinsic activity statistics.** a) Bin spikes of each recorded neuron in each experimental session into 100ms windows to obtain spike-count vectors. b) Perform 5 fold cross-validated latent factor analyses (LFA) with increasing number of factors (from 1 to 25). Determine log-likelihood as a function of the number of factors. Select number of factors (#factors) as the minimal one that yields a relative increment of the cross-validation curve lower than 5%. c) Perform LFA with #factors to split total activity into shared and intrinsic activity components. d) Compute covariance of shared and intrinsic components. e) Compute distribution of covariances for shared and intrinsic components. f) Subsample intrinsic neural activity and compute standard deviation of cross-covariances $\delta c$ as a function of samples (#bins) used. g) Fit dependence of $\delta c$ on the number of bins to estimate and remove bias induced by the finite duration of the recording (see Methods). This procedure yielded an unbiased estimate of the statistics of intrinsic auto- and cross-covariances.

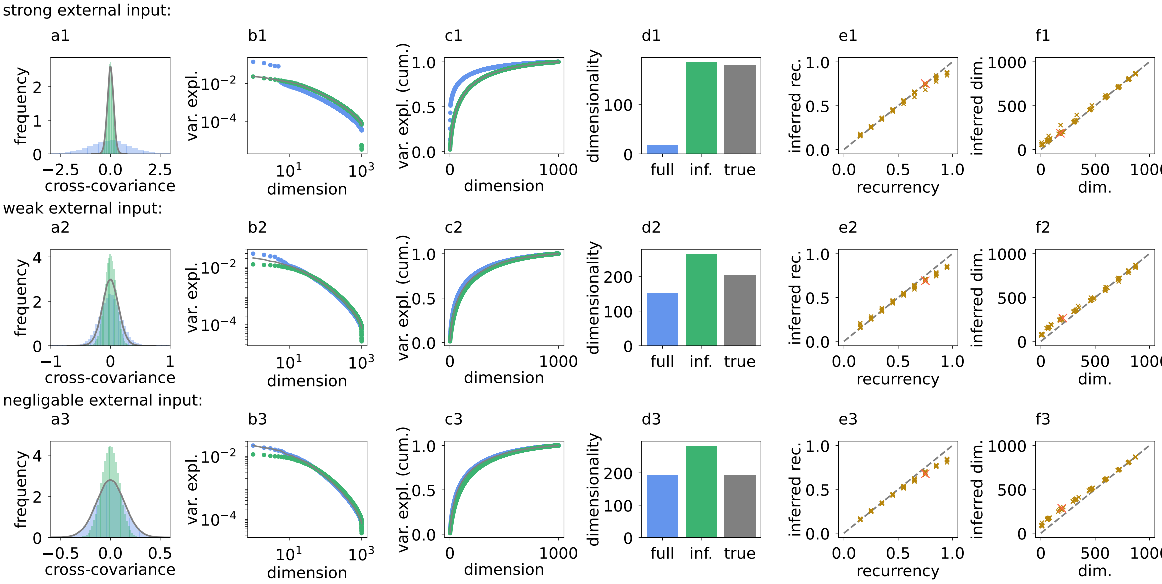

Figure S4: **Latent Factor Analysis applied to simulation data.** LFA yields an upper bound on the dimensionality of intrinsic network activity in linear rate networks. Correlated external inputs can in principle strongly alter covariances within the network, but LFA can extract intrinsic covariances over a wide range of recurrencies (spectral radii) and ranks of external input covariances. In the strongly recurrent regime, our LFA procedure tends to excessively (incorrectly) subtract strong intrinsic activity components, leading to conservative results on recurrency and intrinsic dimensionality. Top row: strong external input ($d_{\text{ext}} = d_{\text{int}}$). Middle row: weak external input ($d_{\text{ext}} = 0.1 d_{\text{int}}$). Bottom row: negligible external input ($d_{\text{ext}} = 0.01 d_{\text{int}}$). a) A low rank input can lead to broader distributions of covariances (blue) with respect to intrinsically generated covariances (gray). Applying LFA, inferred intrinsic covariances (green) recover features of ground-truth intrinsic covariances well. Parameters: recurrency $R = 0.75$, external input rank$(D_{ext}) = 5$. b) Covariance spectra corresponding to distributions in a. c) Cumulative variance explained corresponding to distributions in a. d) Dimensionality of full, inferred intrinsic and true intrinsic activity corresponding to distributions in a. e) Inferred recurrency vs ground truth recurrency for networks of different recurrency $R$ and different ranks of external input (between 1 and 20). Red marker shows configuration shown in a-d. f) Inferred intrinsic dimensionality vs ground truth intrinsic dimensionality for networks of different recurrency $R$ and different ranks of external input (between 1 and 20). More details: see Sec. 4.5.

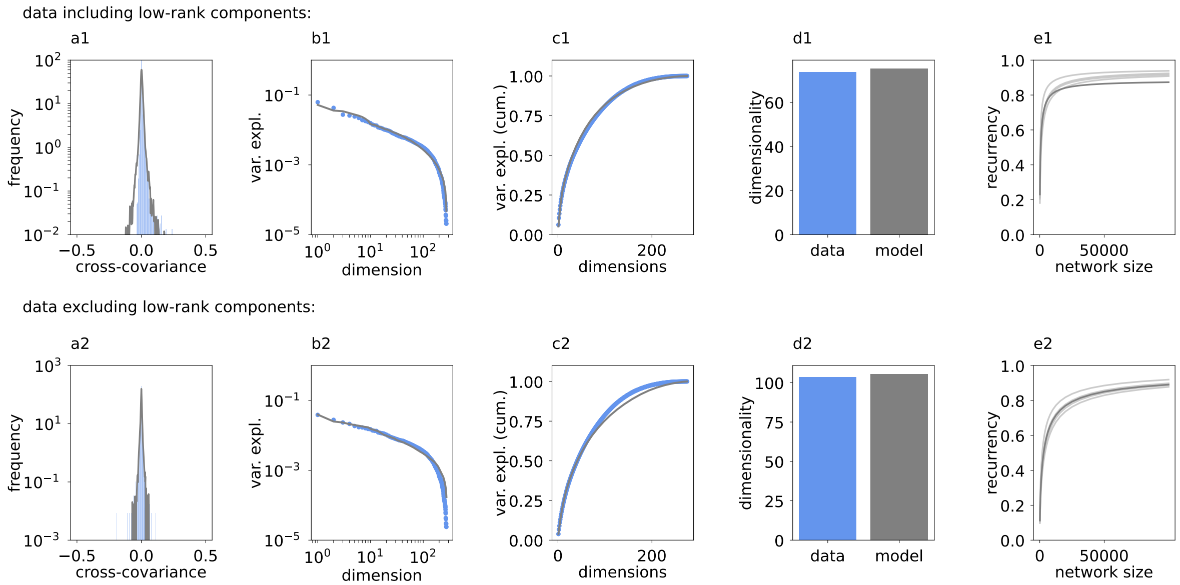

Figure S5: **Fitting of linear network model to mean and standard deviation of auto- and cross-covariances of HMM states in visual cortex.** A linear rate model with Gaussian white noise of variable strength across neurons captures many features of experimentally observed covariances and dimensionality. Top row: fit to data of one example HMM state without removal of low-rank activity components. Bottom row: fit to data of the same HMM state after removal of low-rank activity components using LFA. Even if all removed low-rank activity components were attributed to correlated external inputs and only the activity after LFA is considered to be intrinsic, for reasonable network sizes the network still needs to operate in the strongly recurrent regime to reproduce the data. a) Distribution of experimentally observed covariances (blue) and model fit (gray) in logarithmic frequency scale. b) Variance explained of individual dimensions for data and model fit (rescaled and ranked PCA eigenvalues). c) Cumulative variance explained of data and model fit. d) Dimensionality (participation ratio) of data and model fit. e Inferred recurrency of the fit model as a function of the network size. Dark gray curve shows example HMM state shown in panels a - d. Light gray curves show other HMM states of example session. Parameters: Panels a - d show results of model fits with network size $N = 275$, which equals the number of recorded neurons in the example session. More details: see Sec. 4.6.

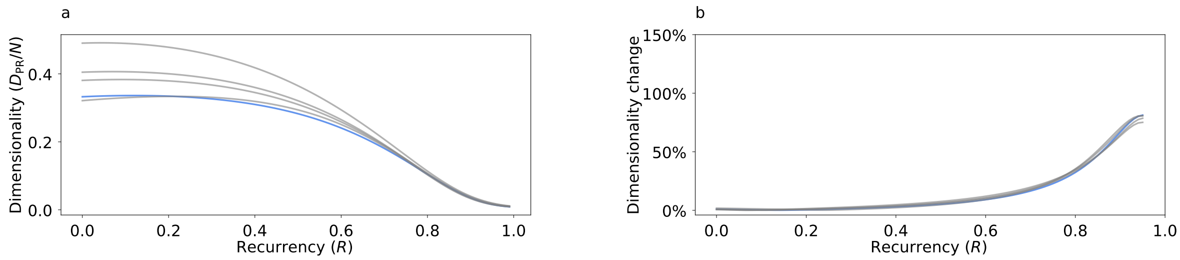

Figure S6: **Dimensionality and its sensitivity to recurrency in the presence of external inputs.** This figure is analogous to Fig. 2c,d of the main text, but with external inputs. Correlated external inputs reduce the dimensionality of network activity, but the overall dependence on and sensitivity to recurrency $R$ is similar to the case without external inputs. a) Dimensionality (normalized to the number of recorded neurons) of activity (including low-rank components) as a function of recurrency for network models from Fig. S5 that were fit to HMM states in visual cortex. Along the x-axis, only the recurrency of the fit model is changed, while all other network and input parameters are preserved. b) Relative dimensionality change as a function of recurrency for $\Delta R = -0.05$ (cf. Fig. 2d in main text) for fit network models in a. More details: see Sec. 4.6.

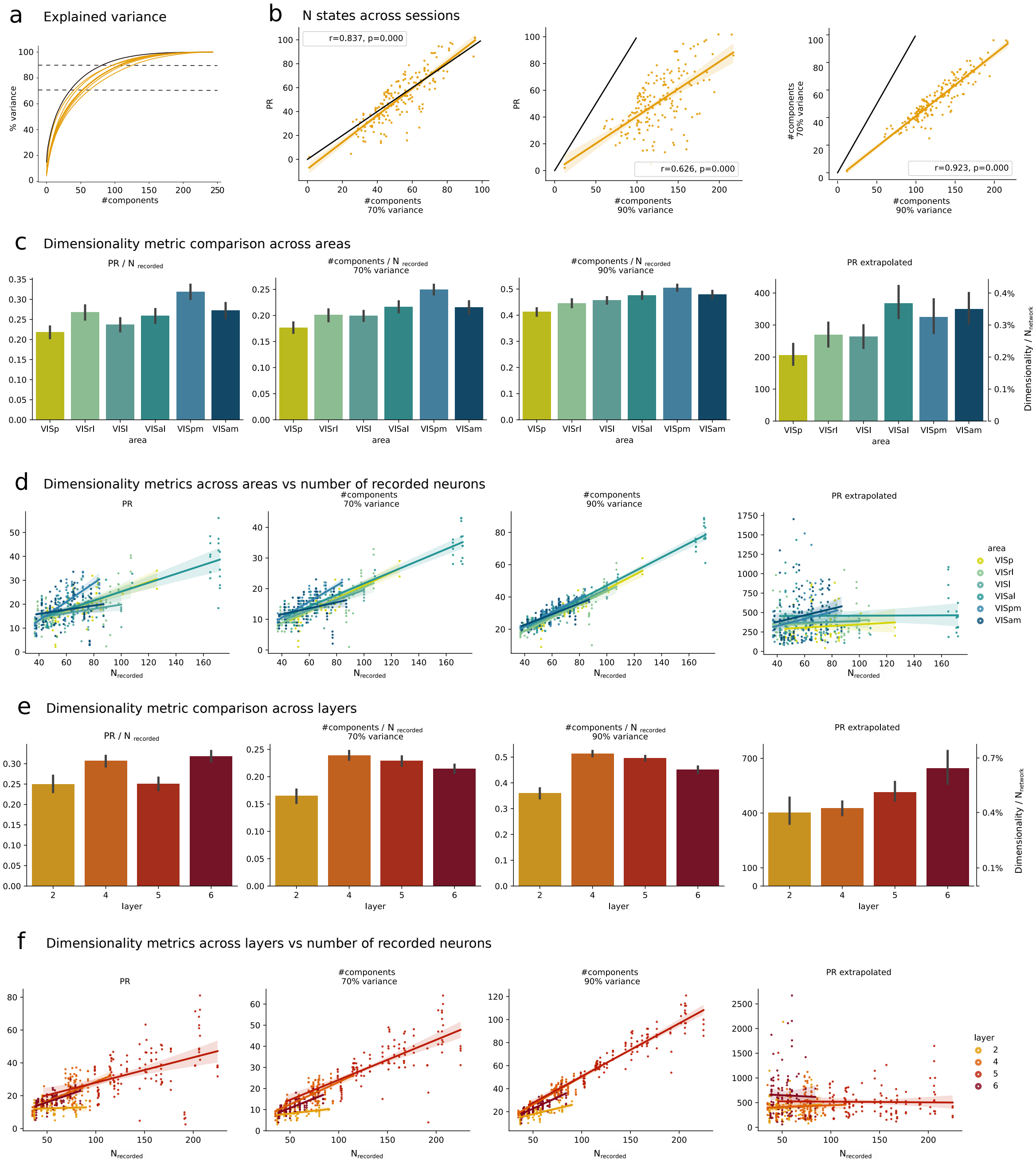

Figure S7: **Dimensionality comparison**. a) Curves of variance explained for different HMM states in one example session. Dashed lines indicate thresholds of 70% and 90% variance explained. The participation ratio $D_{\text{PR}} = 1/\sum_i \tilde{\lambda}_i^2$ depends on all values of the curve (rescaled covariance eigenvalues $\tilde{\lambda}_i = \lambda_i/\sum_k \lambda_k$). b) Left: Participation ratio vs number of components required to explain 70% variance. Center: Participation ratio vs number of components required to explain 90% variance. Right: Number of components required to explain 70% variance vs number of components required to explain 90% variance. c) Comparison of area-resolved dimensionality metrics. First three plots display the same metrics introduced in panel b. The rightmost plot shows the area-resolved extrapolated dimensionality (Eq. S1), also normalized by the assumed network size ($N = 10^5$) on the right y–axis. d) Dependence of dimensionality metrics on the number of recorded neurons (area resolved). e) Same as panel c realized across layers. f) Same as panel d realized across layers.

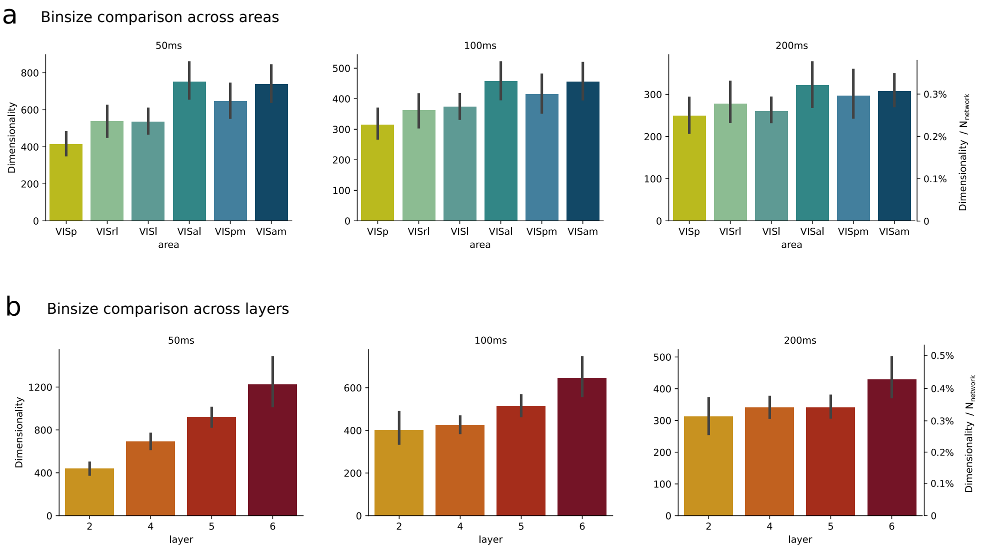

Figure S8: **Dependence of dimensionality on bin size.** a) Dimensionality of activity in different visual areas for bin sizes of 50ms (left), 100ms (middle) and 200ms (right). b) Dimensionality of activity in different cortical layers for bin sizes of 50ms (left), 100ms (middle) and 200ms (right).

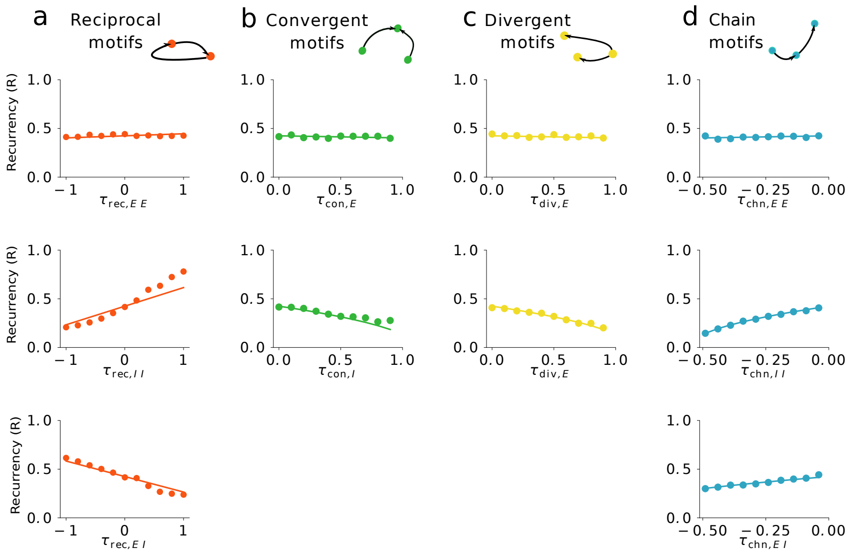

Figure S9: **Numerical validation of theory for recurrency (spectral radius) in excitatory-inhibitory networks with population-specific motif abundances.** Recurrency (spectral radius) $R$ as a function of abundances of excitatory and inhibitory a) recurrent motifs, b) convergent motifs, c) divergent motifs, and d) chain motifs. Only one parameter is varied at a time with all other motif abundances set to zero. Theoretical predictions are given by solid curves, numerical evaluations of single network realizations are shown by colored markers. Cf. Sec. 4.11 for network construction algorithms. The approximate theory correctly predicts the influence of changes in abundance of each type of motif on the spectral radius. In particular, an over-representation of reciprocal EE and II motifs increases the spectral radius while an over-representation of reciprocal EI motifs decreases the spectral radius. These effects are the dominant contributions to the results obtained from the synaptic physiology dataset. More details: see Sec. 4.9ff.

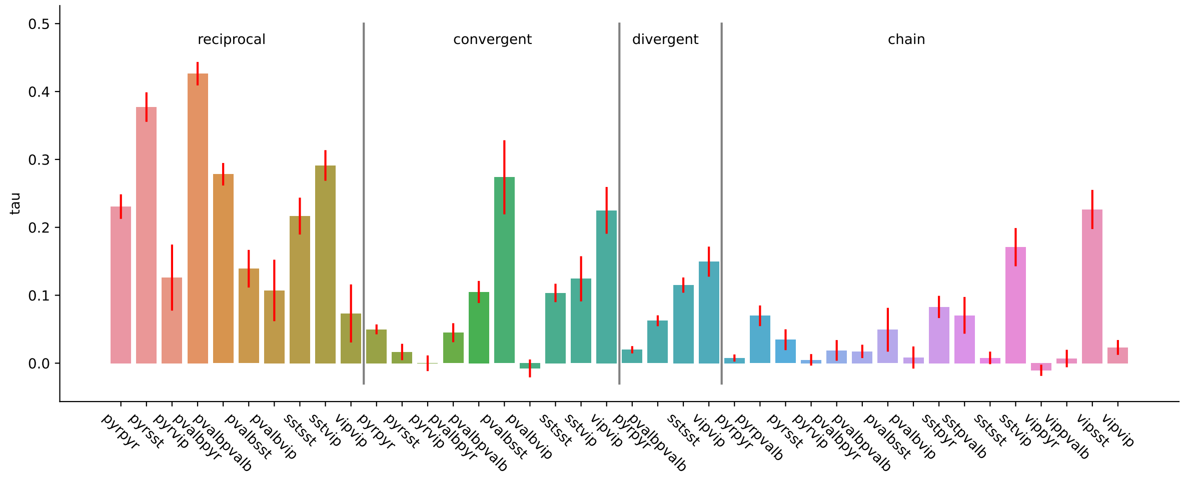

Figure S10: **Motifs of pyramidal neurons with different inhibitory cell types.** Note that there are strong differences in motif abundances among cell types. This points towards differential effects of modulating the activity of different cell-types on network recurrency.

### 3 Methods

#### 3.1 Electrophysiology data and analysis.

##### 3.1.1 Data description summary.

Data are publicly available and a detailed description has been made available by the Allen Institute for Brain Science [1, 2]. The Allen Institute for Brain Science Institutional Animal Care and Use Committee authorized guidelines for mice kept in the animal facility. To identify genetically defined inhibitory cell types via optotagging, most experiments used C57BL/6J wild-type mice ($n = 30$), supplemented by recordings in three transgenic lines ($n = 8$ Pvalb-IRES-Cre x Ai32, $n = 12$ Sst, and $n = 8$ Vip). After surgery, all mice were single-housed and kept on a reverse 12-h light cycle in a shared facility at $20 - 22°$C and 30–70% humidity. Experiments were performed during the dark cycle. Mice had free access to food and water throughout the experiments.

Each mouse had a grade 5 titanium headframe with a cranial window to facilitate co-registration across surgical, intrinsic signal imaging, and electrophysiology rigs. The headframe was glued to a black acrylic photopolymer well that shielded the craniotomy and probes during the experiment and provided a surface for precisely aligning the insertion window.

Mice were mildly anesthetized with 1–1.4% isoflurane and monitored with a PhysioSuite (model PS-MSTAT-RT). Eye drops (Lacri99 Lube Lubricant Eye Ointment; Refresh) kept eyes hydrated and clear during anesthesia. The stimulus we analyzed was a mean brightness grey background and was part of a longer stimulus train (see [2] for details). Mice spent two weeks habituating in sound-attenuated training boxes with headframe holders, running wheels, and stimulus monitors.

Recordings contained 58 experimental sessions in adult mice. At the beginning of the experimental session the cranial coverslip was replaced with an insertion window containing holes aligned to six cortical visual areas. Neural recordings were performed with six Neuropixels probes each containing 960 recording sites providing a maximum of 3.84 mm of tissue coverage. Visual stimuli were generated using scripts based on PsychoPy and followed one of two stimulus sequences ("brain observatory 1.1" and "functional connectivity"). We focused on spontaneous neural activity registered while the animal was not performing any task. In each session the spontaneous activity condition lasted 30 minutes while the animal was in front of a screen of mean grey luminance. We therefore analyzed 26 of the original 58 sessions corresponding to the "functional connectivity" subdataset as they included such period of spontaneous activity. To perform the analysis we used and extended the Allen SDK toolbox https://github.com/AllenInstitute/AllenSDK. The fit and analysis of Hidden Markov Model (HMM) states was performed on the neural activity binned into 5ms windows, while remaining analysis, such as the computation of dimensionality, was performed on neural activity binned into 100ms; see further details below (Fig. S3). Across all analyses we only used those recordings where at least 30 neurons for a specific brain region or brain area were simultaneously recorded.

In addition, layer labels for each cortical unit (L1, L2/3, L4, L5, or L6) were assigned to each neuron using CCFv3 coordinates. The relative thickness of each layer, which can vary both within and between areas, is based on the average of the 1,675 individual brains used to generate the volume template [2].

Cre-lines mice underwent optotagging. Three light stimuli optotagged specific neurons. We determined which units were driven by the optotagging stimulus and likely Cre+ by aligning spikes to these stimuli. Retinal input may have caused spikes after 40 ms, therefore we used the shortest stimulus of 10 ms pulses. These pulses are long enough to observe light-evoked spikes without being tainted by visually induced activity. We identified units that dependably fired faster during the 10 ms pulse across all stimuli. This was done by verifying that the firing rate during the 10ms pulse (2ms to 9ms from stimulus onset) was twice higher than the baseline (-10ms to -2ms from stimulus onset). Further details and code are available (https://allensdk.readthedocs.io/en/latest/_static/examples/nb/ecephys_optotagging.html).

###### 3.1.2 Hidden Markov Model analysis.

The Hidden Markov Model (HMM) analysis of spontaneous activity followed a two stage procedure and was performed using the ssm toolbox (https://github.com/lindermanlab/ssm). Firstly, we binned spike trains into 5ms intervals across the 30 minutes of neural recordings for each session. Then we estimated the number of hidden states using a 5-fold cross-validation procedure. We fit a cross-validated HMM and computed the log-likelihood of fit for an increasing number of hidden states (from 2 to 21). Using the kneed toolkit (https://github.com/arvkevi/kneed), we chose the number of states by means of detecting an elbow in the average log-likelihood across number of states. This algorithm selects a cross-validation curve point on the basis of a single parameter, $s$. We utilized the default value $s = 1$. Using the resulting number of states, we then fit an HMM to the spike counts $n$-times ($n = 5$) and selected the fit with maximum log-likelihood across these. This is to guarantee that the final fit is among the best possible for the selected number of states. The output of the HMM analysis was a confidence (a posterior probability between 0-100%) for each of the underlying hidden factors to have generated the neural activity in each bin. We thresholded this confidence (to 80%) in order to select only temporal intervals in which the algorithm identified a specific hidden factor as the cause of the collective neural activity. Once all time points, and consequently spike-count population vectors, were assigned to one or no hidden state (if no state passed the confidence threshold), we used all such vectors assigned to the same state to calculate a state-specific covariance capturing neural variability for each state. To compute the covariance matrix of neural population vectors, our reference bin size is 100ms, whereas the HMM analysis was performed on 5ms windows of activity. Before calculating the covariance matrix, we therefore rebinned neural activity within each state into 100ms windows, discarding all appearances of an individual state that lasted less than 100ms.

###### 3.1.3 Dimensionality analysis.

We analyzed the participation ratio $D_{\mathrm{PR}}$ [3] as a measure of dimensionality (Fig. S2). This measure can be rewritten in terms of the means $\bar{a}$ and $\bar{c}$ and standard deviations $\delta a$ and $\delta c$ of auto- and cross-covariances, respectively (Fig. S2), and the number of neurons recorded $N_{\mathrm{rec}}$ (similar to [4]):

$$D_{\mathrm{PR}} = \frac{N_{\mathrm{rec}}}{1 + \left(\frac{\delta a}{\bar{a}}\right)^2 + (N_{\mathrm{rec}} - 1)\left(\left(\frac{\delta c}{\bar{a}}\right)^2 + \left(\frac{\bar{c}}{\bar{a}}\right)^2\right)}. \tag{S1}$$

For model data where we had access to all neuron activities, $N_{\mathrm{rec}}$ corresponded to the full network size $N$ (cf. Suppl.Mat. Sec. 4.1 and Sec. 4.2). Applied to experimental data, the dimensionality $D_{\mathrm{PR}}$ depends on the number of recorded neurons. In the absence of any bias in the subsampling procedure the statistics of covariances, as extracted by means of our analysis, are invariant (cf. Suppl.Mat. Sec. 4.2 and Fig. S2) and Eq. S1 is adopted to extrapolate the dimensionality as a function of the neurons recorded $N_{\mathrm{rec}} \to N$.

###### 3.1.4 Bias correction in the statistics of covariances.

We performed a theoretical analysis of the bias on the covariance statistics induced by subsampling both neurons or trials (cf. Suppl.Mat. Sec. 4.3 and [4] for a similar analysis): the empirical estimates $\hat{\bar{a}}$ and $\hat{\bar{c}}$ of the average auto- and cross-covariances are unbiased while the variances $\widehat{\delta a^2}$ and $\widehat{\delta c^2}$ of both auto- and cross-covariances have a bias which decays with the number of trials $N_T$ as $\sim \frac{1}{N_T}$:

$$\bar{a} = \hat{\bar{a}} \tag{S2}$$

$$\bar{c} = \hat{\bar{c}} \tag{S3}$$

$$\delta a^2 = \frac{N_T - 1}{N_T + 1}\widehat{\delta a^2} - \frac{2\left(\hat{\bar{a}}^2 - \hat{\bar{c}}^2\right)}{N_T + 1} + \frac{2\delta c^2}{N_T + 1} \tag{S4}$$

$$\delta c^2 = \frac{N_T - 1}{N_T}\widehat{\delta c^2} - \frac{\hat{\bar{a}}^2 - \hat{\bar{c}}^2}{N_T} - \frac{4}{N + 1}\,\frac{\hat{\bar{c}}^2 - \hat{\bar{a}}\,\hat{\bar{c}}}{N_T} \tag{S5}$$

Here $\hat{}$ indicates the empirical estimate and the non-hat quantities indicate the true values. Based on such analysis we performed a bias correction (cf. Fig. S3 and next section).

###### 3.1.5 Intrinsic covariance analysis.

Under a linear assumption the covariance matrix of neural activity splits into two contributions: a shared and an intrinsic component (cf. Suppl. Mat. Sec. 4.5). In order to estimate these two components we developed a three stage procedure that could be performed by utilizing different algorithms at its core. Here we use Latent Factor Analysis (LFA). In the following we will explain this procedure with LFA but it would work equivalently with PCA or other algorithms. The first stage bins the spikes of neurons into spike counts within non-overlapping windows. We used 100ms bins, Fig. S3a. Then we performed LFA multiple times with an increasing number of hidden factors and computed the log-likelihood as a function of factors with a 5-fold cross-validation technique, Fig. S3b. We selected the number of factors by choosing the corresponding point in the log-likelihood curve where the cross-validated log-likelihood didn't increase more than 5% for the first time. This yielded a robust estimation of where the plateau or peak in the curve is found, Fig. S3b. The second stage estimated the activity of the shared neural activity and intrinsic neural activity by running LFA with the selected number of components, Fig. S3c. The computed shared and intrinsic covariance (Fig. S3d) yielded a first estimate of the standard deviation of intrinsic cross-covariances $\delta c$, Fig. S3e. In the third stage we removed the bias on such estimates by subsampling the intrinsic neural activity (Fig. S3f) and computing $\delta c$ as a function of the number of samples used $N_T$ (Fig. S3g). We then fit the dependence (cf. Eq. S5) $\hat{\delta c}^2 = \delta c^2 + \frac{\text{const.}}{N_T}$ to extract the true value of $\delta c$ from the estimates $\hat{\delta c}$. All analyses were run through custom scripts based on the scikit learn library.

###### 3.1.6 Dimensionality analysis and HMM states

The HMM analysis mapped intervals in the spontaneous activity to a number of hidden latent states whose appearance often corresponded to changes in the behavior of the animal (Figs. 1c to 1d). We then compared the covariance statistics obtained separately in each state to the covariance statistics obtained in the entire interval of spontaneous activity (Figs. 1i to 1l). Also, we compared these two conditions to the one where for each neural state we performed a cross-validated Latent Factor Analysis (LFA) over the covariance statistics. This removed any remaining non-stationary component potentially not detected by the HMM.

In Figs. 2e to 2f we computed the dimensionality change across layers and brain areas. This is defined as $\Delta D_{\text{PR}} = \frac{2|D_{\text{PR}}(A) - D_{\text{PR}}(B)|}{D_{\text{PR}}(A) + D_{\text{PR}}(B)}$ where $A, B$ are two separate conditions with distinct activity statistics. In all these analyses these two conditions correspond to a different neural state. In each recording session we took all possible combinations of two HMM states. We computed $D_{\text{PR}}$ in each of the two arbitrarily assigning one state to condition $A$, and the second to condition $B$ (notice that $\Delta D_{\text{PR}}$ is symmetric in $A, B$). We then computed $\Delta D_{\text{PR}}$ averaging across all states and recording sessions. The only difference across panels and bars in Fig. 2e is that the computation of dimensionality was restricted to only those neurons of a given layer or brain area. In the control condition we split the statistics of states into two parts, the first and second half, and utilized these two as conditions $A, B$.

In Fig. 2f we performed a similar analysis. We computed the dimensionality change across all neurons between any two states. Then we restricted the statistics to only neurons of a given cell type, computing their firing rate difference across the two states. In order to correlate the two quantities, dimensionality change and firing rate difference, we analyzed a modified version of $\Delta D_{\text{PR}} = \frac{2(D_{\text{PR}}(A) - D_{\text{PR}}(B))}{D_{\text{PR}}(A) + D_{\text{PR}}(B)}$ (notice the absence of a modulus in the numerator). This was correlated with $\Delta_{\text{FR}} = \text{FR}(A) - \text{FR}(B)$, where $\text{FR}(A), \text{FR}(A)$ denote the average firing rate in the two conditions.

#### 3.2 Synaptic physiology data and analysis.

##### 3.2.1 Data description summary

We analyzed recently published data collected at the Allen Institute for Brain Science [5], where synaptic connections were probed using simultaneous patch clamp of 3-to-8 cell groups. The dataset reports the synapses (or lack thereof) between any pair of probed neurons. Over 32000 synaptic connections were probed resulting in 1368 chemical synapses from mouse primary visual cortex and 363 from human cortex.

Multiple methodologies were utilized to identify each subclass of cell based on cell type and layer position. Brightfield, epifluorescence (tdTomato and EGFP) and dye-filled recording pipettes were used to acquire images of the recording site. By combining epifluorescence and pipette dye, transgenic expressions in cells were then identified. The genetic identification of each cell was complemented or supplemented by a morphological examination (see [5] for details).

We analyzed the data in two ways: (1) accounting for the cell-type of pre- and postsynaptic neurons (pyramidal cells Pyr, somatostatin cells SST, vasoactive intestinal polypeptide cells VIP, parvalbumin cells PV), as well as the excitatory or inhibitory nature of the synapses, and (2) ignoring these divisions into cell types, but still accounting for the excitatory or inhibitory nature of the synapses.

##### 3.2.2 Extraction of connection probabilities and motif statistics from synaptic physiology data

We first describe the analysis in Figs. 4a to 4e. In this analysis we considered all synapses and first report their statistics across either layers (Fig. 4b) or cell types (Fig. 4c). Based on these statistics we then computed the value of $\sigma$ as in ($R = \sigma \cdot R_{\text{motifs}}$) via the multi-population analysis introduced in [6] and detailed for our application in Supp. Mat. Sec. 4.7. To compute the relative sigma, as displayed in Figs. 4d to 4e, we first computed a baseline value $\sigma_{baseline}$. In the layer-resolved analysis, this baseline corresponds to the full circuit of excitatory and inhibitory populations across layers 2 and 5. We then computed the relative decrease in $\sigma$ upon limiting the analysis to the subcircuit of excitatory and inhibitory populations of a given layer Fig. 4d. In the cell-type specific analysis, the baseline considered synapses of all inhibitory cell types to be equally strong. In this case we computed the relative increase or decrease in $\sigma$ upon strengthening the efficacy of synapses of a given cell type (multiplying their strength by a factor 2, Fig. 4e), for details see Supp. Mat. Sec. 4.7).

We then proceeded to compute the motif statistics as shown in Figs. 4f to 4m. To estimate $R_{\text{motifs}}$ from the whole dataset or a subset of data (bootstraps) we proceeded as follows: Each motif type considered in this work consists of two connections. In our theoretical work, we for simplicity only keep track of the presynaptic cell type of each connection (see Supp. Mat. Sec. 4.9). Divergent motifs are thus computed per presynaptic cell type, irrespective of the two postsynaptic cell types. Convergent motifs take into account the identity of both presynaptic cells. As the postsynaptic neuron of one connection in a reciprocal motif is the presynaptic neuron of the other connection, our analysis includes the identity of pre- and postsynaptic neurons. For chain motifs, we take into account both presynaptic neurons which implies that only the final neuron of the chain is left unspecified. To obtain consistency between our theory and data analysis, we therefore also calculated motif probabilities from the data on this level of granularity.

We first computed the probabilities $p_\alpha$ of having an excitatory ($\alpha = E$) or inhibitory ($\alpha = I$) synapse among two neurons and estimated the variances $\sigma_\alpha^2 = p_\alpha(1 - p\alpha)$ according to Bernoulli statistics. In addition, for reciprocal and chain motifs, we calculated probabilities $p_{\beta\alpha}$ of having a connection between a presynaptic neuron of type $\alpha$ and postsynaptic neuron of type $\beta$, along with the respective variances. These probabilities (first order statistics) are required to normalize the motif probabilities (second order statistics) to determine whether there is an over- or underrepresentation of motifs compared to an assumption of independent connectivity statistics.

We then computed the probabilities of having a reciprocal, chain, convergent or divergent motif in the data: In each session where $n$ (= 8 or fewer) neurons were probed we computed the total number of pairs (reciprocal motifs) or triplets of neurons (divergent, convergent, chain motifs) that could carry a corresponding motif. For example for $n = 8$ excitatory neurons the number of possible excitatory reciprocal motifs is $8 \times 7/2 = 28$. We then counted the number of motifs in each session of each type, taking into account the identity of presynaptic neurons as described above. Finally we summed up all the occurrences for each motif type and the total number of pairs or triplets of neurons that could potentially carry such motif. The division between these two numbers yielded the probability of occurrence for each motif. We then subtract the independent first order statistics $p_\alpha^2$ in the case of divergent motifs $(\alpha \to (X, X))$, $p_\alpha p_\beta$ for convergent motifs $((\alpha, \beta) \to X)$, $p_{\alpha\beta} p_{\beta\alpha}$ for reciprocal motifs $(\alpha \leftrightarrow \beta)$ and $p_\beta p_{\beta\alpha}$ for chain motifs $(\alpha \to \beta \to X)$ and divide by the respective $\sigma$'s to obtain correlation coefficients $\tau_{\text{rec}}, \tau_{\text{chn}}, \tau_{\text{div}}, \tau_{\text{con}}$ for the individual populations (Fig. 4f). Error bars in Fig. 4f show one standard deviation across bootstraps that take into account 80% of the data. We further combined excitatory and inhibitory motif statistics into effective $\tau$'s as described in Suppl.Mat. Sec. 4.9 taking into account an estimate of the effective relative strength $g$ between inhibitory and excitatory synapses and the relative population size $\gamma$. Finally, we applied the formula in Eq. 3 to obtain the recurrency $R$ (Figs. 4g to 4i).

For Fig. 4h, to compare the empirical distribution to a null hypothesis, we generated ensembles of 8 neurons (as many as simultaneously probed in the dataset) and randomly defined each neuron as either excitatory or inhibitory. We randomly generated inhibitory or excitatory synapses, with probabilities matching the probability of having excitatory or inhibitory synapses in the dataset. On this "random" dataset we then performed the same analysis as on the empirical dataset: we subsampled the entire statistics 500 times. Each of these subsamples contained 80% of the total statistics, that is 80% of all sessions chosen randomly. Finally we computed $R_{\text{motifs}}$ for each of these, Fig. 4h.

For the analysis in Figs. 4j to 4m we performed the same procedure as described above for cell types (Figs. 4j to 4k) and layers (Figs. 4l to 4m). For cell types we strengthened synapses whose presynaptic site was of a given cell type (by a factor 2). For layers we limited our analyses to the excitatory and inhibitory cells of that specific layer.

Analyses on human data were performed with the same procedures.

##### 3.2.3 Reanalysis of synaptic physiology data from Song et al.

We analyzed data from [7] on motif abundances in excitatory connections of rat visual cortex and evaluated their influence on $R_{\text{motifs}}$ in an analogous way as we did for the synaptic physiology dataset [5, 8]. The probability of occurrence for each motif type is given by the ratio between the observed occurrences and the possible occurrences of motifs, each summed across all simultaneous patch clamp recordings of triplets of neurons. The data in [7] is presented in their Fig. 4b in terms of 16 different patterns which each contain possibly multiple of the motifs discussed here. For example, pattern 12 contains one reciprocal motif, one divergent motif, two convergent motifs and two chain motifs. We thus calculated the probability of each motif in terms of the occurrences $M_p$ of patterns $p \in \{1, 2, ..., 16\}$, given by the numbers above the bars in Fig. 4b of [7]), and the number of occurrences of this particular motif in each pattern:

$$p_{\text{motif}} = \frac{\sum_{\text{triplets}} \text{observed motifs}}{\sum_{\text{triplets}} \text{possible motifs}} = \frac{\sum_{p=1}^{16} M_p \cdot \text{motifs in p}}{\sum_{p=1}^{16} M_p \cdot \text{possible motifs in triplet}} . \tag{S6}$$

This yielded values $\tau_{\text{rec}} = 0.40, \tau_{\text{chn}} = 0.02, \tau_{\text{div}} = 0.02, \tau_{\text{con}} = 0.04$ and $R_{\text{motifs}} = 1.34$, which is broadly consistent with our findings on excitatory connections from the synaptic physiology dataset [5, 8].

#### 3.3 Theoretical and computational analysis

##### 3.3.1 Network models and linear response theory.

We made use of the fact that correlations in spontaneous, asynchronous irregular activity states of spiking networks can be well understood using linear response theory [9, 10]: starting from a network of leaky integrate-and-fire (LIF) neurons, linearization around some stationary state maps the statistics of fluctuations to an equivalent set of Ornstein-Uhlenbeck processes coupled via some effective connectivity matrix [11]. The stationarity required for linearization holds if population-level fluctuations are not strongly amplified by large excess excitation. In our analysis of experimental data we ensure stationarity by studying covariances and dimensionality within individual HMM states.

Ornstein-Uhlenbeck processes are linear stochastic differential equations that can be analyzed using statistical field theory [12, 13]. Fig. 2c shows that such theory faithfully predicts the statistics of covariances and dimensionality as a function of the recurrency in direct simulations of LIF neurons. The recurrency $R$ is a summary measure of the network's effective connectivity and formally corresponds to the spectral radius of bulk connectivity eigenvalues (for details see Supp. Mat. Sec. 4.8). Our theoretical derivations first focused on networks with homogeneous connection statistics. We then expanded this theoretical analysis to investigate networks with excitatory and (multiple) inhibitory populations and distinct connection statistics for the different types of neurons (Supp. Mat. Sec. 4.9). Our results linking the dimensionality and the spectral radius also generalize well to more complex network topologies (Fig. 2c).

##### 3.3.2 Theory of spectral radius in networks with second order motifs.

Using the path-integral representation of coupled Ornstein-Uhlenbeck processes [13], we performed an average of the moment-generating function for the network dynamics over the statistics of connections. Second-order connection motifs were thereby incorporated via the covariance tensor $\Delta_{ijkl} = \langle W_{ik} W_{jl} \rangle - \langle W_{ik} \rangle \langle W_{jl} \rangle$ between connections $W_{ik}$ from neuron $k$ to neuron $i$ and $W_{jl}$ from neuron $l$ to neuron $j$. Similar to the case of reciprocal connections [14, 15], the second-order connectivity statistics yield non-Gaussian integrals that cannot be solved exactly. We obtained good approximations to these integrals for large networks by using a saddle-point approximation of auxiliary fields that were introduced for the terms related to the various motif contributions. The associated self-consistency equations for the saddle points showed parameter-dependent divergence structures that we related to connectivity eigenvalues crossing the line of instability of the linear network. By distinguishing between outlier and bulk eigenvalues, this analysis allowed us to infer a theoretical prediction of the spectral radius of the effective connectivity in relation to the various motif abundances, cf. Suppl. Mat. Sec. 4.8 and Sec. 4.9.

##### 3.3.3 Numerical validation of motifs theory.

Following [16] we numerically generated network connectivity matrices as a superposition of Gaussian i.i.d. matrices

$$W_{ij} = \mu + A\nu_{ij} + B\nu_{ji} + C\eta_i + D\eta_j \, . \tag{S7}$$

In Suppl. Mat. Sec. 4.11 we show that coefficients $A, B, C, D$ can be chosen such that the second order statistics of $W$ reflects the correlations induced by various motif abundances in sparse networks. Note that this Gaussian connectivity does not contain any higher-order cumulant information of networks with motifs. The effect of such higher-order cumulants is, however, typically suppressed by the large network size of biological networks, as discussed in Suppl. Mat. Sec. 4.8. It turns out that Eq. S7 cannot generate any arbitrary constellation of motif abundances, as detailed in Suppl. Mat. Sec. 4.11. In particular, correlations induced by divergent and convergent motifs are typically positive, and chain motif correlations cannot be generated independently from other motifs. These considerations give rise to the $\tau$ ranges shown in Fig. 3, where we validated our theoretical predictions for the spectral radius and dimensionality. The spectral radius is well predicted for all values of $\tau$ (Fig. 3b). The same holds true for the prediction of the dimensionality (Fig. 3c), except for reciprocal motifs, where the prediction is only correct on a qualitative level. The prediction of the dimensionality relies - in addition to our results on the motif dependence of the spectral radius - on the mapping between the spectral radius and the width the covariance distribution that has been derived in [12] for homogeneous random networks using beyond-mean-field techniques. This relation is robust as long as eigenvalue spectra of connectivities show a circular organization in the complex plane. Convergent, divergent and chain motifs do not strongly alter the shape of the bulk of connectivity eigenvalues [17]. Therefore, the theory for homogeneous random networks yields correct quantitative results for these cases. Reciprocal motifs, however, deform the bulk eigenvalues from the circular to an elliptic shape [18], which causes the minor quantitative mismatch between theory and simulations.

### 4 Theoretical Supplementary Material

#### 4.1 Statistics of covariances and dimensionality

The dimensionality of network dynamics is defined as the participation ratio

$$D_{\mathrm{PR}} = \frac{1}{\sum_{i=1}^{N} \tilde{\lambda}_i^2} \tag{S8}$$

with normalized eigenvalues $\tilde{\lambda}_i = \lambda_i / \sum_k \lambda_k$ of the covariance matrix $C$ between neurons. The eigenvalues of the covariance matrix are related to the trace of the matrix, such that, for a network of $N$ neurons, the dimensionality of the full network dynamics can be rewritten as [4]

$$D_{\mathrm{PR}} = \frac{tr(C)^2}{tr(C^2)} = \frac{N}{1 + \left(\frac{\delta a}{\bar{a}}\right)^2 + (N-1)\left(\left(\frac{\delta c}{\bar{a}}\right)^2 + \left(\frac{\bar{c}}{\bar{a}}\right)^2\right)}. \tag{S9}$$

#### 4.2 Subsampling and dimensionality extrapolation

In experiments one has access to only a subset of neurons of a given neural network. The dimensionality $D_{\mathrm{PR,rec}}$ of the recorded parallel activity is given by

$$D_{\mathrm{PR,rec}} = \frac{N_{\mathrm{rec}}}{1 + \left(\frac{\delta a}{\bar{a}}\right)^2 + (N_{\mathrm{rec}}-1)\left(\left(\frac{\delta c}{\bar{a}}\right)^2 + \left(\frac{\bar{c}}{\bar{a}}\right)^2\right)}, \tag{S10}$$

which is formally identical to (S9) but with the network size $N$ being replaced by the number of recorded neurons $N_{\mathrm{rec}}$. The dimensionality $D_{\mathrm{PR,rec}}$ of the recorded activity therefore depends on the number of recorded neurons rather than the network size. Furthermore, the statistics of covariances is based on the subsampled covariance matrix. In the absence of any bias in the subsampling procedure the statistics of covariances are invariant, meaning that one can recover the dimensionality $D_{\mathrm{PR}}$ of the full network dynamics by replacing $N_{\mathrm{rec}}$ with the assumed underlying network size $\to N$ in (S10), thus extrapolating to this assumed size, as shown in Fig. S2f and Fig. 1, Fig. 2. We note that for spatially structured networks a local recording using e.g. a Neuropixels probe, yields an estimate for covariances that is only valid for the local network surrounding the electrodes. Extrapolation of dimensionality far beyond such local network sizes is therefore not valid. Keeping the statistics of covariances fixed during extrapolation yields a saturating dimensionality for $N_{\mathrm{rec}} \to \infty$. Note further that estimators for variances of auto- and cross-covariances are biased when covariances are based on a finite number of trials. In this case, the estimators also receive a correction that depends on the number of recorded neurons, which can, however, be removed as noted above and in Sec. 4.3).

#### 4.3 Bias correction in statistics of covariances

For experimental data, covariances are estimated based on a limited number of trials. In consequence, the estimators for the dispersion of auto- and cross-covariances may be biased towards larger variances. To find analytical correction formulas for this finite data bias we assume a Gaussian probability distribution $p(n^1, ..., n^{N_T})$, $n^k \in \mathbb{N}_0^N$, of activities $n_i^k$ of neuron $i$ in trial $k \in \{1, ..., N_T\}$. We further assume that there are no correlations between different trials, so that the probability distribution factorizes over trials and the moment generating function is

$$\phi(l^1, ..., l^{N_T}) = \prod_{k=1}^{N_T} \exp\left(m^{\mathrm{T}} l^k + \frac{1}{2} l^{k,\mathrm{T}} c l^k\right), \tag{S11}$$

where $m$ is the vector of mean activities, $c$ is the covariance matrix with the following meta-statistics: mean autocovariance $\bar{a}$, variance of autocovariances $\delta a^2$, mean cross-covariance $\bar{c}$ and variance of cross-covariances $\delta c^2$ across neurons. Since we are only interested in the statistics of covariances, we can, without loss of generality, set all mean activities to zero

$$\phi(l^1, ..., l^{N_T}) \to \prod_i \exp\left(c_{ii}\frac{1}{2}\sum_{k=1}^{N_T} l_i^k l_i^k\right)\prod_{i<j}\exp\left(c_{ij}\sum_{k=1}^{N_T} l_i^k l_j^k\right) \tag{S12}$$

The meta-statistics are assumed to be identical across trials. Furthermore, we assume the meta-statistics to be the same for all realizations of distributions of covariances across neurons. In the following, $\langle\rangle$ denotes the average over these realizations obtained from the averaged moment generating function

$$\begin{aligned}
\left\langle \phi(l^1, ..., l^{N_T}) \right\rangle &= e^{\bar{\Phi}(l^1, ..., l^{N_T})} \\
\bar{\Phi}(l^1, ..., l^{N_T}) &= \frac{1}{2}\bar{a}\sum_i \sum_{k=1}^{N_T} l_i^k l_i^k + \frac{1}{8}\delta a^2 \sum_i \sum_{k,l=1}^{N_T} l_i^k l_i^k l_i^l l_i^l \\
&+ \frac{1}{2}\bar{c}\sum_{i\neq j}\sum_{k=1}^{N_T} l_i^k l_j^k + \frac{1}{4}\delta c^2 \sum_{i\neq j}\sum_{k,l=1}^{N_T} l_i^k l_j^k l_i^l l_j^l .
\end{aligned}$$

Therefore we obtain second and fourth order cumulants of activities

$$\begin{aligned}
\langle n_a^m n_b^n \rangle = \langle\langle n_a^m n_b^n \rangle\rangle &= \left.\frac{\partial^2 \bar{\Phi}}{\partial l_a^m \partial l_b^n}\right|_{l=0} \\
&= \delta_{mn}\left(\delta_{ab}\bar{a} + (1-\delta_{ab})\bar{c}\right) \; , \\
\langle\langle n_a^m n_b^n n_c^o n_d^p \rangle\rangle &= \left.\frac{\partial^4 \bar{\Phi}}{\partial l_a^m \partial l_b^n \partial l_c^o \partial l_d^p}\right|_{l=0} \\
&= \delta a^2 \delta_{ab}\delta_{ac}\delta_{ad}\left(\delta_{mn}\delta_{op} + \delta_{mo}\delta_{np} + \delta_{mp}\delta_{no}\right) \\
&+ \delta c^2 \delta_{ac}\delta_{bd}\left(1-\delta_{ab}\right)\left(\delta_{mn}\delta_{op} + \delta_{mp}\delta_{no}\right) \\
&+ \delta c^2 \delta_{ab}\delta_{cd}\left(1-\delta_{ac}\right)\left(\delta_{mo}\delta_{np} + \delta_{mp}\delta_{no}\right) \\
&+ \delta c^2 \delta_{ad}\delta_{bc}\left(1-\delta_{ab}\right)\left(\delta_{mn}\delta_{op} + \delta_{mo}\delta_{np}\right) .
\end{aligned}$$

Let $\hat{\cdot}$ denote the empirical estimates of mean activities and covariances from $N$ recorded neurons in $N_T$ trials of the experiment. Unbiased estimators for sample means and covariances are (using Bessel's correction)

$$\begin{aligned}
\hat{m}_i &= \frac{1}{N_T}\sum_{k=1}^{N_T} n_i^k \; , \\
\hat{c}_{ij} &= \frac{1}{N_T - 1}\sum_{k=1}^{N_T} (n_i^k - \hat{m}_i)(n_j^k - \hat{m}_j) \; .
\end{aligned}$$

The latter can be expressed purely in terms of activities

$$\hat{c}_{ij} = \sum_{k,l=1}^{N_T}\left(\frac{\delta_{kl}}{N_T} - \frac{1-\delta_{kl}}{N_T(N_T-1)}\right) n_i^k n_j^l .$$

These estimators are unbiased because

$$\langle \hat{m}_i \rangle = \frac{1}{N_T} \sum_{k=1}^{N_T} \underbrace{\langle n_i^k \rangle}_{\bar{m}} = \bar{m} = 0\,,$$

$$\begin{aligned}\langle \hat{c}_{ij} \rangle &= \sum_{k,l=1}^{N_T} \left( \frac{\delta_{kl}}{N_T} - \frac{1-\delta_{kl}}{N_T(N_T-1)} \right) \langle n_i^k n_j^l \rangle \\ &= \sum_{k,l=1}^{N_T} \left( \frac{\delta_{kl}}{N_T} - \frac{1-\delta_{kl}}{N_T(N_T-1)} \right) \delta_{kl} \left( \delta_{ij}\overline{a} + (1-\delta_{ij})\overline{c} \right) \\ &= \delta_{ij}\overline{a} + (1-\delta_{ij})\overline{c}\ .\end{aligned}$$

The mean covariance defined as

$$\hat{\overline{a}} = \frac{1}{N} \sum_{i=1}^{N} \hat{c}_{ii}\,, \tag{S13}$$

$$\hat{\overline{c}} = \frac{1}{N(N-1)} \sum_{i \neq j}^{N} \hat{c}_{ij} \tag{S14}$$

is therefore also unbiased

$$\langle \hat{\overline{a}} \rangle = \frac{1}{N} \sum_{i=1}^{N} \langle \hat{c}_{ii} \rangle = \overline{a}\,, \tag{S15}$$

$$\langle \hat{\overline{c}} \rangle = \frac{1}{N(N-1)} \sum_{i \neq j}^{N} \langle \hat{c}_{ij} \rangle = \overline{c}\,. \tag{S16}$$

The variance of auto- and cross-covariances needs to be properly defined using Bessel's correction

$$\begin{aligned}\delta\hat{a}^2 &= \frac{1}{N-1} \sum_{a=1}^{N} \left( \hat{c}_{aa} - \hat{\overline{a}} \right)^2 \\ &= \sum_{a,b}^{N} \left( \frac{\delta_{ab}}{N} - \frac{1-\delta_{ab}}{N(N-1)} \right) \hat{c}_{aa}\hat{c}_{bb}\,, \\ \delta\hat{c}^2 &= \frac{1}{N(N-1)-2} \sum_{a \neq b}^{N} \left( \hat{c}_{ab} - \hat{\overline{c}} \right)^2 \\ &= \frac{1}{N(N-1)} \sum_{a,b \neq}^{N} \hat{c}_{ab}\hat{c}_{ab} \\ &\quad - 4 \frac{1}{N(N-1)-2} \frac{1}{N(N-1)} \sum_{a,b,d \neq}^{N} \hat{c}_{ab}\hat{c}_{ad} \\ &\quad - \frac{1}{N(N-1)-2} \frac{1}{N(N-1)} \sum_{a,b,c,d \neq}^{N} \hat{c}_{ab}\hat{c}_{cd}\,.\end{aligned}$$

Note that Bessel's correction amounts to a normalization $\frac{1}{N-1}$ for the variances (subtract one degree of freedom as usual) and $\frac{1}{N(N-1)-2}$ for cross-covariances (subtract two degrees of freedom due to symmetry $\hat{c}_{ab} = \hat{c}_{ba}$). These corrections are the right ones as can be seen by calculating the estimated variances of variances

$$\left\langle \hat{\delta a^2} \right\rangle = \sum_{a,b}^{N} \left( \frac{\delta_{ab}}{N} - \frac{1-\delta_{ab}}{N(N-1)} \right) \langle \hat{c}_{aa} \hat{c}_{bb} \rangle$$
$$= \delta a^2 + \frac{2\left(\delta a^2 - \delta c^2\right)}{N_T - 1} + \frac{2\left(\overline{a}^2 - \overline{c}^2\right)}{N_T - 1}$$

and cross-covariances

$$\left\langle \hat{\delta c^2} \right\rangle = \frac{1}{N(N-1)} \sum_{a,b\neq}^{N} \langle \hat{c}_{ab} \hat{c}_{ab} \rangle$$
$$- 4 \frac{1}{N(N-1)-2} \frac{1}{N(N-1)} \sum_{a,b,d\neq}^{N} \langle \hat{c}_{ab} \hat{c}_{ad} \rangle$$
$$- \frac{1}{N(N-1)-2} \frac{1}{N(N-1)} \sum_{a,b,c,d\neq}^{N} \langle \hat{c}_{ab} \hat{c}_{cd} \rangle$$
$$= \delta c^2 + \frac{\delta c^2 + \overline{a}^2 - \overline{c}^2}{N_T - 1} + \frac{4}{N+1} \frac{\overline{c}^2 - \overline{ac}}{N_T - 1},$$

where the true variances $\delta a^2$ and $\delta c^2$ on the right hand sides of the equations are not receiving any correction due to the number of recorded neurons $N$. To obtain the above expressions, we used

$$\langle \hat{c}_{ab} \hat{c}_{cd} \rangle = \sum_{m,n,o,p=1}^{N_T} \left( \frac{\delta_{mn}}{N_T} - \frac{1-\delta_{mn}}{N_T(N_T-1)} \right) \left( \frac{\delta_{op}}{N_T} - \frac{1-\delta_{op}}{N_T(N_T-1)} \right) \langle n_a^m n_b^n n_c^o n_d^p \rangle$$
$$= \delta a^2 \delta_{ab} \delta_{ac} \delta_{ad} \left( 1 + \frac{2}{N_T - 1} \right)$$
$$+ \delta c^2 \delta_{ac} \delta_{bd} \left(1 - \delta_{ab}\right) \left( 1 + \frac{1}{N_T - 1} \right)$$
$$+ \delta c^2 \delta_{ab} \delta_{cd} \left(1 - \delta_{ac}\right) \frac{2}{N_T - 1}$$
$$+ \delta c^2 \delta_{ad} \delta_{bc} \left(1 - \delta_{ab}\right) \left( 1 + \frac{1}{N_T - 1} \right)$$
$$+ \left(\delta_{ab} \overline{a} + (1-\delta_{ab}) \overline{c}\right) \left(\delta_{cd} \overline{a} + (1-\delta_{cd}) \overline{c}\right)$$
$$+ \frac{1}{N_T - 1} \left(\delta_{ad} \overline{a} + (1-\delta_{ad}) \overline{c}\right) \left(\delta_{bc} \overline{a} + (1-\delta_{bc}) \overline{c}\right)$$
$$+ \frac{1}{N_T - 1} \left(\delta_{ac} \overline{a} + (1-\delta_{ac}) \overline{c}\right) \left(\delta_{bd} \overline{a} + (1-\delta_{bd}) \overline{c}\right)$$

and

$$\left\langle n_i^m n_j^n n_k^o n_l^p \right\rangle = \left\langle \left\langle n_i^m n_j^n n_k^o n_l^p \right\rangle \right\rangle + \left\langle n_i^m n_j^n \right\rangle \left\langle n_k^o n_l^p \right\rangle + \left\langle n_i^m n_l^p \right\rangle \left\langle n_j^n n_k^o \right\rangle + \left\langle n_i^m n_k^o \right\rangle \left\langle n_j^n n_l^p \right\rangle . \quad \text{(S17)}$$

In summary, these results provide a procedure to infer the true statistics of covariances from estimators based on $N$ recorded neurons in $N_T$ trials:

$$
\begin{aligned}
\bar{a} &= \hat{\bar{a}}\,,\\
\bar{c} &= \hat{\bar{c}}\,,\\
\delta a^2 &= \frac{N_T-1}{N_T+1}\delta\hat{a}^2 - \frac{2\left(\hat{\bar{a}}^2-\hat{\bar{c}}^2\right)}{N_T+1} + \frac{2\delta c^2}{N_T+1}\,,\\
\delta c^2 &= \frac{N_T-1}{N_T}\delta\hat{c}^2 - \frac{\hat{\bar{a}}^2-\hat{\bar{c}}^2}{N_T} - \frac{4}{N+1}\frac{\hat{\bar{c}}^2-\hat{\bar{a}}\hat{\bar{c}}}{N_T}\,,
\end{aligned}
$$

with

$$
\begin{aligned}
\hat{\bar{a}} &= \frac{1}{N}\sum_{i=1}^{N}\hat{c}_{ii}\,,\\
\hat{\bar{c}} &= \frac{1}{N(N-1)}\sum_{i\neq j}^{N}\hat{c}_{ij}\,,\\
\delta\hat{a}^2 &= \frac{1}{N-1}\sum_{a=1}^{N}\left(\hat{c}_{aa}-\hat{\bar{a}}\right)^2\,,\\
\delta\hat{c}^2 &= \frac{1}{N(N-1)-2}\sum_{a\neq b}^{N}\left(\hat{c}_{ab}-\hat{\bar{c}}\right)^2\,.
\end{aligned}
$$

#### 4.4 Relation between intrinsic covariances and network recurrency

The participation ratio Eq. S9 formally depends on all first and second moments of auto- and cross-covariances. Yet, due to the large prefactor $N-1$ in the denominator, mainly cross-neuronal coordination in form of the rescaled mean $m=\frac{\bar{c}}{\bar{a}}$ and standard deviation $s=\frac{\delta c}{\bar{a}}$ of cross-covariances determine the dimensionality. For intrinsic covariances we found that $s$ is much larger than $m$ (Fig. 1k), so that in good approximation the intrinsic dimensionality is determined by the variability of intrinsic cross-covariances across neuron pairs:

$$\frac{D_{\mathrm{PR}}}{N} = \frac{1}{1+Ns^2}\,. \tag{S18}$$

Intrinsic correlations in networks have been explained using linear response theory [19, 20, 21, 9, 10]. Linearization around some stationary state yields an effective connectivity that couples neuronal fluctuations around their mean activities [20, 22]. Such fluctuations can be formally described with a "linear rate model," a coupled set of Ornstein-Uhlenbeck processes [23]

$$\tau_i\partial_t x_i(t) = -x_i(t) + \sum_{j=1}^{N} W_{ij}x_j(t) + \xi(t)\,, \tag{S19}$$

where the centered Gaussian white noise $\xi(t)$ reflects intrinsic stochasticity of neuronal processes. Its variance $\left\langle \xi(t)\xi(s)^{\mathrm{T}}\right\rangle = \delta(t-s)D$ derives from the mean activity of neurons in the stationary state that is used as a working point for linearization [11].

Reference [12] used beyond-mean field techniques to study the statistics of correlations in such systems and found that, in homogeneous networks, the width $s$ of the covariance distribution is determined by the network recurrency $R$, that is the spectral radius of the effective connectivity

$$Ns^2 = \frac{1}{\left(1-R^2\right)^2} - 1. \tag{S20}$$

Inserting (S20) into (S18) yields a relation between the dimensionality of the full network dynamics and the recurrency of the network, via the relationship

$$\frac{D_{\mathrm{PR}}}{N} = (1 - R^2)^2. \tag{S21}$$

The robustness of equation (S20) with respect to other network topologies shown in [12] implies a robustness of equation (S21) (cf. Fig. 2c of the main text). To investigate the relation between dimensionality and connectivity, it is therefore sufficient to study how the recurrency of the network, i.e. the spectral radius of the effective connectivity, is determined by the network topology (see Suppl. Mat. S4.8-4.9).

##### 4.4.1 Effect of other covariance statistics on dimensionality

Apart from the contribution of $s$ to the dimensionality, one can employ the generalized results on the statistics of auto- and cross-covariances derived in [24] to refine how the dimensionality depends on network parameters. More specifically, for a network with mean connectivity $\langle W_{ij} \rangle = \mu$ and variance $\langle W_{ij}^2 \rangle - \langle W_{ij} \rangle^2 = \sigma^2/N$ as well as intrinsic stochasticity of mean $\langle D_{ii} \rangle = \bar{D}$ and variance $\langle D_{ii}^2 \rangle - \langle D_{ii} \rangle^2 = \overline{\delta D^2}$ one obtains

$$\bar{a} = \left(1 + 2M + NM^2\right) \frac{\bar{D}}{1 - R^2} \tag{S22}$$

$$\bar{c} = \left(2M + NM^2\right) \frac{\bar{D}}{1 - R^2} \tag{S23}$$

and

$$\overline{\delta c^2} = \left(2M^2\right) \overline{\delta D^2} + \bar{R}\frac{1}{N}(1 + 2M)^2 \overline{\delta D^2} + \bar{R}\left\{\left(2(1 + 2M)M^2 + NM^4\right) \overline{\delta D^2}\right\} + \bar{R}\frac{1}{N}\left(\bar{a}^2 + (N - 1)\bar{c}^2\right) \tag{S24}$$

$$\overline{\delta a^2} = (1 + 4M)\overline{\delta D^2} + 2\overline{\delta c^2} \tag{S25}$$

with

$$M = \frac{\mu}{1 - N\mu}$$
$$R = (1 + 2M + M^2)\sigma^2$$
$$\bar{R} = 2\frac{R^2}{1 - R^2} + \left(\frac{R^2}{1 - R^2}\right)^2$$

Note that these equations reduce to the results in [12] for $M \to 0$ and $\overline{\delta D^2} = 0$. The above expressions can be inserted in Eq. S9 to obtain a more detailed dependence of dimensionality on the network parameters.

#### 4.5 Extraction of intrinsic covariances from driven network dynamics using Latent Factor Analysis

In this section we apply the latent factor analysis (LFA) performed on the experimental data (Fig. S3) to model data, from an input-driven network model with known ground truth intrinsic activity and external drive. To construct the model data we make use of the linear rate model Eq. S19 introduced above.

For networks that receive correlated external inputs, the mean of those inputs changes the firing rate as well as the gain of neurons, and thereby the effective connectivity $W$ [24]. Temporal fluctuations of external inputs directly drive local correlations. This latter effect can be modeled in the framework above as an additional independent, centered and correlated external input $\eta(t)$, which we here assume to have a covariance $\langle \eta(t)\eta(s)^{\mathrm{T}} \rangle = \delta(t-s)D_{\mathrm{external}}$.

We are interested in covariances of spike counts computed for large time bins. These covariances are equivalent to time-lag integrated covariances or zero-frequency cross-spectra [12]. For the network dynamics Eq. S19 including the additional external drive $\eta$, we obtain these zero-frequency cross-spectra by studying the zero-frequency Fourier mode $X$ of the dynamics

$$0 = -X + WX + \tilde{\xi} + \tilde{\eta} \tag{S26}$$

where $\tilde{\xi}$ and $\tilde{\eta}$ are the zero-frequency components of the inputs $\xi(t)$ and $\eta(t)$, respectively. $\tilde{\xi}$ and $\tilde{\eta}$ follow Gaussian statistics with zero mean and covariance $D_{\mathrm{intrinsic}}$ and $D_{\mathrm{external}}$, respectively. Eq. (S26) can be solved for $X$, such that

$$X = (1-W)^{-1}(\tilde{\xi} + \tilde{\eta}). \tag{S27}$$

The zero-frequency mode $X$ therefore follows Gaussian statistics

$$X \sim \mathcal{N}(0, C) \tag{S28}$$

with

$$C = C_{\mathrm{intrinsic}} + C_{\mathrm{external}} \tag{S29}$$

$$C_{\mathrm{intrinsic}} = (1-W)^{-1} D_{\mathrm{intrinsic}} (1-W^{\mathrm{T}})^{-1} \tag{S30}$$

$$C_{\mathrm{external}} = (1-W)^{-1} D_{\mathrm{external}} (1-W^{\mathrm{T}})^{-1}. \tag{S31}$$

The theory developed in this study linking dimensionality and network recurrency, see Sec. S4.8-4.9, strictly holds for intrinsic network activity, i.e. $D_{\mathrm{external}} = 0$. In the experiments we analyzed, however, local recorded networks are potentially driven by external inputs as well. Such external inputs may affect local cross-covariances and the dimensionality. This effect could even persist after removing strong activity transients by restricting the analysis to individual HMM states. In order to still apply our theory, we need to first extract the intrinsic covariances $C_{\mathrm{intrinsic}}$ from the measured covariances $C$ by removing the externally induced components $C_{\mathrm{external}}$.

The extraction of $C_{\mathrm{intrinsic}}$ is based on the following conservative assumption: all low-rank activity inferred by Latent Factor Analysis [25] is attributed to external inputs. Latent Factor Analysis uses an expectation maximization procedure to infer low-dimensional latent factors $\tilde{X}$ in the activity $X$ such that the covariance

$$C_{LFA} = C_{\mathrm{shared}} + R$$

can be split into a shared low-rank component $C_{\mathrm{shared}} = \langle X_{\mathrm{shared}} X_{\mathrm{shared}}^{\mathrm{T}} \rangle$ and a diagonal remainder $R$. Here, $X_{\mathrm{shared}}$ denotes the projection of the low-dimensional latent factors into the high-dimensional space of neurons using the loading matrix $L$: $X_{\mathrm{shared}} = L\tilde{X}$. In case of an ideal fit, $C = C_{LFA}$ and latent factors have zero mean and unit covariance such that the shared covariance reduces to $C_{\mathrm{shared}} = LL^T$. However, since the above splitting procedure cannot be done exactly for every covariance matrix $C$, the above operation is in general an approximation. Therefore we introduce the covariance $\Delta C$ as the mismatch between $C$ and $C_{LFA}$, and write

$$\begin{aligned} C &= C_{\mathrm{shared}} + R + \Delta C \\ &=: C_{\mathrm{shared}} + \hat{C}_{\mathrm{intrinsic}}. \end{aligned}$$

Assuming that the external component $C_{\mathrm{external}}$ is the only low-rank component of $C$, LFA yields $C_{\mathrm{shared}} = C_{\mathrm{external}}$ such that $\hat{C}_{\mathrm{intrinsic}} = C_{\mathrm{intrinsic}}$. In this case, we can extract an estimate of $C_{\mathrm{intrinsic}}$ as

$$\hat{C}_{\mathrm{intrinsic}} = C - C_{\mathrm{shared}}\,, \tag{S32}$$

where $C = \langle XX^{\mathrm{T}} \rangle$ and $C_{\mathrm{shared}} = \langle X_{\mathrm{shared}} X_{\mathrm{shared}}^{\mathrm{T}} \rangle$.

Fig. S4 shows a systematic analysis of the above procedure for the case of strong ($\langle\eta_i^2\rangle = \langle\xi_i^2\rangle$), weak ($\langle\eta_i^2\rangle = 0.1\langle\xi_i^2\rangle$) and vanishingly small ($\langle\eta_i^2\rangle = 0.01\langle\xi_i^2\rangle$) external input. For all these input scenarios, the LFA procedure yields conservative estimates for the intrinsic covariances, as well as the recurrent coupling and dimensionality: Strong external inputs significantly broaden the distribution of cross-covariances (Fig. S4a) and change the spectrum of covariance eigenvalues for low PC dimensions (Fig. S4b). As a result, the dimensionality is reduced (Fig. S4c,Fig. S4d). Both effects can be removed by extracting $\hat{C}_{\mathrm{intrinsic}}$ via a latent factor analysis. This works well for low and intermediate spectral radii, as can be seen from the correct inference of the recurrency and dimensionality in these regimes (Fig. S4e,Fig. S4f). For large spectral radii, the dimensionality of $C_{\mathrm{intrinsic}}$ is low, so that $C_{\mathrm{shared}}$ of LFA not only extracts $C_{\mathrm{external}}$ but also components from $C_{\mathrm{intrinsic}}$. The vanishingly small input case thereby serves as a control, because almost all inferred low-rank activity is not due to external input but rather due to intrinsic activity. The results for this control case and the other input cases show that the LFA procedure with parameters chosen as for the experimental data underestimates recurrency and overestimates intrinsic dimensionality. Therefore LFA yields conservative results in terms of how strongly connectivity constrains dimensionality.

We deliberately chose a rather simple model setting for the parameter scan in the previous results. In particular, low-rank external inputs were projected into the network with random loadings drawn from a Gaussian distribution. One consequence of this simple choice of external input is for example the gap in the spectrum of eigenvalues in Fig. S4b that is typically not present in experimental data. In the next section, we show that linear models can be fit to better reproduce the experimentally observed covariance features.

#### 4.6 Fitting linear network models to experimental data

Eq. S22-Eq. S25 relate the mean and variance of intrinsic auto- and cross-covariances to the five network parameters $\mu, \sigma, \bar{D}, \overline{\delta D^2}$ and $N$, which are the mean and standard deviation of connections, the mean and variance of intrinsic neuron stochasticity $D_{\mathrm{intrinsic}}$ and the network size, respectively. For every network size $N$, we can invert these relations and find network parameters to reproduce the given means and variances of covariance in the experimental data. Although the fit is only based on the four statistics of covariances, Fig. S5 shows that the model can largely reproduce the curves of variance explained and many features of the covariance distribution. In the top row, where all activity is assumed to be generated intrinsically (no correlated external input), a few strong covariances cannot be captured by the model (Fig. S5a1); this, however, does not have a strong impact on the shape of the variance explained curves (Fig. S5b,Fig. S5c). These unpredicted strong covariances get subtracted using the LFA procedure, which is shown in the bottom row where low-rank activity components have been removed using LFA (Fig. S5a2). In this latter case, we assumed that all low-rank activity stems from external input. The remaining activity can also be fit well by the linear model. In both scenarios (top: all activity is treated as intrinsic, bottom: only activity after LFA subtraction is treated as intrinsic), the linear network models need to operate in the strongly recurrent regime to reproduce the data (Fig. S5e).

In Fig. S6 we analyze how the dimensionality of network activity changes with recurrency in the presence of external input. To do this for plausible external inputs we follow a five-step procedure: 1) We perform LFA to split the covariance of a given HMM state into $C_{\mathrm{intrinsic}}$ and $C_{\mathrm{external}}$. 2) We fit network parameters $\mu, \sigma, \bar{D}, \overline{\delta D^2}$ on the mean and variances of auto- and cross-covariances $C_{\mathrm{intrinsic}}$ (cf. Fig. S5 bottom). For the fitting, we choose the network size to be equal to the number of recorded neurons in the data. 3) We construct a realization of a Gaussian connectivity matrix $W$ and a Gaussian noise matrix $D_{\mathrm{intrinsic}}$ based on the inferred network parameters. 4) We infer the external input covariance $D_{\mathrm{external}}$ based on $C_{\mathrm{external}}$ obtained in step 1 and the connectivity matrix $W$ obtained in step 3 via $D_{\mathrm{external}} = (1 - W)C_{\mathrm{external}}(1 - W^T)$. 5) We rescale the connectivity to modify the recurrency of the model to a desired value leaving all other network and input parameters ($\mu, D_{\mathrm{intrinsic}}, D_{\mathrm{external}}$) the same.

The resulting change in dimensionality with recurrency for fixed external input is shown in Fig. S6 as an average over 50 repetitions of the procedure (realizations of $W$). The external input overall reduces the dimensionality, but the general trend with which dimensionality is further reduced with stronger recurrency is similar to the case without inputs. Also similarly to the case without inputs, the relative change in dimensionality with respect to a change in recurrency is largest in the strongly recurrent regime.

#### 4.7 Theory of spectral radius in multi-population networks

The result from [6] on the spectral radius $R = \sigma$ of block-structured connectivity matrices is

$$\sigma = \sqrt{\max(\lambda(M))}\,, \tag{S33}$$

where $\lambda(M)$ are eigenvalues of the matrix $M$ that contains the variances of each block of the connectivity and the size of the block:

$$M_{\alpha\beta} = \gamma_\beta \mathcal{G}^2_{\alpha\beta} = N_\beta \frac{\mathcal{G}^2_{\alpha\beta}}{N}\,. \tag{S34}$$

Here $\gamma_\beta = \frac{N_\beta}{N}$ is the fraction of all cells that belong to population $\beta$ and $\frac{\mathcal{G}^2_{\alpha\beta}}{N}$ is the variance of connections in block $(\alpha, \beta)$. For Bernoulli connectivity, the variance is given by the connection probability $p_{\alpha\beta}$ and the synaptic strength $J_{\alpha\beta} = g_{\alpha\beta} J$ of connections from population $\beta$ to population $\alpha$:

$$\frac{\mathcal{G}^2_{\alpha\beta}}{N} = p_{\alpha\beta}(1 - p_{\alpha\beta}) g^2_{\alpha\beta} J^2\,. \tag{S35}$$

So in total one obtains

$$\begin{aligned} M_{\alpha\beta} &= N J^2 \gamma_\beta p_{\alpha\beta}(1 - p_{\alpha\beta}) g^2_{\alpha\beta} \\ &= N J^2 m_{\alpha\beta}\,, \end{aligned}$$

which has an overall scaling factor $NJ^2$ and a block-specific component $m_{\alpha\beta}$ that depends on population sizes, population specific connection probabilities and relative synaptic efficacy factors.

##### 4.7.1 Relative recurrency

In Fig. 4d and Fig. 4e we compare the recurrency of a circuit with the recurrency of modified versions of that circuit. In case of the layer analysis (Fig. 4d), the circuit corresponds to 4 populations (2 excitatory, 2 inhibitory) spread across layers 2 and 5. The modified circuits are the subcircuits within either layer 2 or 5, respectively. For $X \in \{2, 5\}$, we obtain the relative recurrency as

$$\frac{\sigma_{LX}}{\sigma_{all}} = \sqrt{\frac{\max(\lambda(M_{LX}))}{\max(\lambda(M_{all}))}} = \sqrt{\frac{\max(\lambda(m_{LX}))}{\max(\lambda(m_{all}))}}$$

with

$$m_{all} = \begin{pmatrix} m_{L2E,L2E} & m_{L2E,L2I} & m_{L2E,L5E} & m_{L2E,L5I} \\ m_{L2I,L2E} & m_{L2I,L2I} & m_{L2I,L5E} & m_{L2I,L5I} \\ m_{L5E,L2E} & m_{L5E,L2I} & m_{L5E,L5E} & m_{L5E,L5I} \\ m_{L5I,L2E} & m_{L5I,L2I} & m_{L5I,L5E} & m_{L5I,L5I} \end{pmatrix} \tag{S36}$$

and

$$m_{L2} = \begin{pmatrix} m_{L2E,L2E} & m_{L2E,L2I} & 0 & 0 \\ m_{L2I,L2E} & m_{L2I,L2I} & 0 & 0 \\ 0 & 0 & 0 & 0 \\ 0 & 0 & 0 & 0 \end{pmatrix}, \quad m_{L5} = \begin{pmatrix} 0 & 0 & 0 & 0 \\ 0 & 0 & 0 & 0 \\ 0 & 0 & m_{L5E,L5E} & m_{L5E,L5I} \\ 0 & 0 & m_{L5I,L5E} & m_{L5I,L5I} \end{pmatrix}. \tag{S37}$$

In case of the cell-type specific analysis (Fig. 4e), the circuits consist of one pyramidal cell population and three inhibitory populations (PV, SST and VIP). The modified circuits model the case where one of the inhibitory populations has an increased synaptic efficacy with respect to the others, e.g. $g_{\alpha,PV} = 2 \cdot g_{\alpha,SST} = 2 \cdot g_{\alpha,VIP}$ for the case of a PV dominated circuit. The relative recurrency of the modified circuit with strengthened population $\kappa$ is then given by

$$\frac{\sigma_\kappa}{\sigma_{\text{ref}}} = \sqrt{\frac{\max(\lambda(M_\kappa))}{\max(\lambda(M_{\text{ref}}))}} = \sqrt{\frac{\max(\lambda(m_\kappa))}{\max(\lambda(m_{\text{ref}}))}}\,,$$

where $m_{\text{ref}}$ has the reference values $g_{\alpha,PV} = g_{\alpha,SST} = g_{\alpha,VIP}$ and $m_\kappa$ has an increased $g_{\alpha,\kappa}$ by a factor 2.

#### 4.8 Derivation of spectral radius in homogeneous networks with second-order motifs

##### 4.8.1 Moment generating functional and disorder average

The analysis starts from the equation for the moment generating function of the time-averaged activity $\mathbf{X}$ (cf. (S26) for a network of linear coupled rate neurons, [12])

$$Z(\mathbf{J}) = |\det(\mathbf{1}-\mathbf{W})| \int_{\mathbf{X},\tilde{\boldsymbol{X}}} \exp\left(\tilde{\boldsymbol{X}}^T(\mathbf{1}-\mathbf{W})\boldsymbol{X} + \frac{1}{2}\tilde{\mathbf{X}}^T\boldsymbol{D}\tilde{\mathbf{X}} + \mathbf{J}\mathbf{X}\right)$$

In the following, we assume uniform noise covariance $\boldsymbol{D} = D\mathbb{1}$ as the noise amplitude and correlation does not determine stability. Our goal is to calculate the ensemble average $\langle Z(\mathbf{J})\rangle_W$ of the moment generating functional over all realizations of the effective connectivity $\mathbf{W}$. Dahmen et al. [12] showed that the $\mathbf{W}$-dependence of the prefactor $|\det(\mathbf{1}-\mathbf{W})|$ can, to a good approximation, be factorized such that

$$\langle Z(\mathbf{J})\rangle_W \approx \frac{\left\langle \tilde{Z}(\mathbf{J})\right\rangle_W}{\left\langle \tilde{Z}(0)\right\rangle_W}\,,$$

where we defined $\tilde{Z}(\mathbf{J}) = \int_{\mathbf{X},\tilde{\boldsymbol{X}}} \exp\left(\tilde{\boldsymbol{X}}^T(\mathbf{1}-\mathbf{W})\boldsymbol{X} + \frac{D}{2}\tilde{\mathbf{X}}^T\tilde{\mathbf{X}} + \mathbf{J}\mathbf{X}\right)$. To calculate the ensemble average we need to compute the integral $\int D\mathbf{W} = \prod_{i,j}\int \mathrm{d}W_{ij}$ of $\tilde{Z}(\mathbf{J})$ and weigh each contribution $\mathbf{W}$ according to its joint probability $p(\mathbf{W})$ with mean connection strengths $\langle W_{ij}\rangle = \mu_{ij}$ and covariance of connection strengths $\langle\langle W_{ik}W_{jl}\rangle\rangle = \frac{\Delta_{ijkl}}{N}$

$$\begin{aligned}
\left\langle \tilde{Z}(\mathbf{J})\right\rangle_W &= \int_{\mathbf{X},\tilde{\boldsymbol{X}}} \exp\left(-\tilde{\boldsymbol{X}}^T\boldsymbol{X} + \frac{D}{2}\tilde{\mathbf{X}}^T\tilde{\mathbf{X}} + \mathbf{J}\mathbf{X}\right) \\
&\times \int D\mathbf{W}p(\mathbf{W})\exp\left(\tilde{\boldsymbol{X}}^T\mathbf{W}\boldsymbol{X}\right) \\
&\approx \int_{\mathbf{X},\tilde{\boldsymbol{X}}} \exp\left(-\tilde{\boldsymbol{X}}^T\boldsymbol{X} + \frac{D}{2}\tilde{\mathbf{X}}^T\tilde{\mathbf{X}} + \mathbf{J}\mathbf{X}\right) \\
&\times \exp\left(\sum_{i,j}\mu_{ij}\tilde{X}_iX_j + \frac{1}{2N}\sum_{i,j,k,l}\Delta_{ijkl}\tilde{X}_i\tilde{X}_jX_kX_l\right).
\end{aligned}$$

Note that, when applying the often-used scaling of connection strengths $W_{ij} \sim 1/\sqrt{N}$ with network size $N$ [26], we get $\mu_{ij} = \mathcal{O}(1/\sqrt{N})$ and $\Delta_{ijkl} = \mathcal{O}(1)$, and higher-order connectivity cumulants are suppressed, as we assume large networks.

##### 4.8.2 Second order motifs

Second order motifs are encoded in the covariance $\Delta_{ijkl}$ as

$$\begin{aligned}\Delta_{ijkl} &= \langle\langle W_{ik}W_{jl}\rangle\rangle\, N\\ &= \sigma^2\left(\delta_{ij}\delta_{kl}+\tau_{\mathrm{div}}(1-\delta_{ij})\delta_{kl}+\tau_{\mathrm{con}}(1-\delta_{kl})\delta_{ij}\right)\\ &+ \sigma^2\left(\tau_{\mathrm{chn}}\left(\delta_{il}(1-\delta_{jk})+\delta_{jk}(1-\delta_{il})\right)+\tau_{\mathrm{rec}}\delta_{il}\delta_{jk}\right)\,,\end{aligned}$$

such that we obtain a “‘four-point interaction term“‘ of the form

$$\begin{aligned}\frac{1}{\sigma^2}\sum_{i,j,k,l}\Delta_{ijkl}\tilde{X}_i\tilde{X}_jX_kX_l &= \underbrace{(1-\tau_{\mathrm{div}}-\tau_{\mathrm{con}})}_{=:A}\sum_i\tilde{X}_i\tilde{X}_i\sum_k X_kX_k\\ &+ \underbrace{(\tau_{\mathrm{rec}}-2\tau_{\mathrm{chn}})}_{=:B}\left(\sum_i\tilde{X}_iX_i\right)^2\\ &+ \tau_{\mathrm{div}}\left(\sum_i\tilde{X}_i\right)^2\sum_k X_kX_k\\ &+ \tau_{\mathrm{con}}\sum_i\tilde{X}_i\tilde{X}_i\left(\sum_k X_k\right)^2\\ &+ 2\tau_{\mathrm{chn}}\sum_i\tilde{X}_iX_i\left(\sum_{j,k}\tilde{X}_jX_k\right)\,,\end{aligned} \tag{S38}$$

where we used $\sum_{i\neq j}X_iX_j=\left(\sum_i X_i\right)^2-\sum_i X_iX_i$. The first two lines of the above equation show a similar structure as in [18, 27]: inserting a field $Q=A\frac{\sigma^2}{N}\sum_k X_kX_k$ yields a renormalization of the noise amplitude $D$, while the insertion of a field $P=B\frac{\sigma^2}{N}\sum_k\tilde{X}_kX_k$ results in a renormalization of the propagator. The other lines $3-4$ in (S38) contain terms $R=\sum_k X_k$ and $S=\sum_i\tilde{X}_i$ which on expectation vanish exactly. Therefore, we could introduce the auxiliary fields $R$ and $S$ along with $P$ and $Q$, but in the saddle-point approximation, we would find that $R^*=\sum_k\langle X_k\rangle=0$ and $S^*=\sum_i\left\langle\tilde{X}_i\right\rangle=0$ such that, at this level of approximation, we can neglect lines $3-4$ in (S38). We could also express line 5 using $R$ and $S$, which would in principle lead to no contribution. However, line 5 will yield a renormalization of the response propagator, which is informative about outlier eigenvalues of the connectivity that are being induced by chain motifs, as will be discussed below. Introducing auxiliary fields $P,Q$ and $T=\frac{\tau_{\mathrm{chn}}\sigma^2}{N}\sum_{j,k}\tilde{X}_jX_k$ as well as their conjugate variables arising from the Fourier representation of the delta distribution yields

$$\begin{aligned}e^{\frac{A\sigma^2}{2N}\sum_i\tilde{X}_i\tilde{X}_i\sum_k X_kX_k} &\sim \int_{Q,\tilde{Q}} e^{-\frac{N}{A\sigma^2}Q\tilde{Q}+\tilde{Q}\sum_k X_kX_k+\frac{1}{2}Q\sum_i\tilde{X}_i\tilde{X}_i}\,,\\ e^{\frac{B\sigma^2}{2N}\left(\sum_i\tilde{X}_iX_i\right)^2} &\sim \int_P e^{-\frac{N}{2B\sigma^2}P^2+P\sum_i\tilde{X}_iX_i}\,,\\ e^{\frac{\tau_{\mathrm{chn}}\sigma^2}{N}\sum_i\tilde{X}_iX_i\sum_{j,k}\tilde{X}_jX_k} &\sim \int_{T,\tilde{T}} e^{-\frac{N}{\tau_{\mathrm{chn}}\sigma^2}T\tilde{T}+\tilde{T}\sum_{j,k}\tilde{X}_jX_k+T\sum_i\tilde{X}_iX_i}\end{aligned}$$

as well as the disorder-averaged generating function

$$\begin{aligned}\left\langle \tilde{Z}(\mathbf{J})\right\rangle_{W} & \sim \int_{P,Q,\tilde{Q},T,\tilde{T}} e^{-\frac{N}{A\sigma^{2}}Q\tilde{Q}-\frac{N}{2B\sigma^{2}}P^{2}-\frac{N}{\tau_{\mathrm{chn}}\sigma^{2}}T\tilde{T}+\ln\Omega(P,Q,T)},\\ \Omega\left(P,Q,T\right) & = \int_{\mathbf{X},\tilde{\mathbf{X}}} e^{-\tilde{\boldsymbol{X}}^{T}\boldsymbol{K}\boldsymbol{X}+\frac{D+Q}{2}\tilde{\mathbf{X}}^{T}\tilde{\mathbf{X}}+\tilde{Q}\boldsymbol{X}^{T}\boldsymbol{X}+\mathbf{J}\mathbf{X}},\end{aligned}$$

where we define the inverse propagator

$$\boldsymbol{K}:=\Big((1-P-T)\mathbf{1}-\boldsymbol{\mu}-\tilde{T}\{\mathbf{1}\}\Big)$$

with $\mathbf{1}$ denoting the matrix of ones. As for fully random networks [12], we evaluate the exponent of the disorder-averaged generating function in saddle-point approximation, which yields

$$\tilde{Q}^{*}\sim\left\langle \tilde{\mathbf{X}}^{T}\tilde{\mathbf{X}}\right\rangle =0$$

and, in addition,

$$Q^{*} = A\frac{\sigma^{2}}{N}\left\langle \boldsymbol{X}^{T}\boldsymbol{X}\right\rangle_{P^{*},Q^{*},T^{*}} = A\frac{\sigma^{2}}{N}\sum_{i}\left[\boldsymbol{K}^{-1}(D+Q^{*})\boldsymbol{K}^{-T}\right]_{ii}, \tag{S39}$$

$$P^{*} = B\frac{\sigma^{2}}{N}\left\langle \tilde{\boldsymbol{X}}^{T}\boldsymbol{X}\right\rangle_{P^{*},Q^{*},T^{*}} = B\frac{\sigma^{2}}{N}\sum_{i}\left[\boldsymbol{K}^{-1}\right]_{ii}, \tag{S40}$$

$$T^{*} = \tau_{\mathrm{chn}}\frac{\sigma^{2}}{N}\left\langle \tilde{\boldsymbol{X}}^{T}\{\mathbf{1}\}\boldsymbol{X}\right\rangle_{P^{*},Q^{*},T^{*}} = \tau_{\mathrm{chn}}\frac{\sigma^{2}}{N}\sum_{i,j}\left[\boldsymbol{K}^{-1}\right]_{ij}, \tag{S41}$$

$$\tilde{T}^{*} = \tau_{\mathrm{chn}}\frac{\sigma^{2}}{N}\left\langle \tilde{\boldsymbol{X}}^{T}\boldsymbol{X}\right\rangle_{P^{*},Q^{*},T^{*}}. \tag{S42}$$

In the following, we investigate the divergences in these equations as these give insight into the eigenvalues of the connectivity: The network activity becomes unstable when eigenvalues of the propagator diverge. From the propagator for the full linear network, $(\mathbf{1}-\boldsymbol{W})^{-1}$, this occurs when eigenvalues of $\boldsymbol{W}$ have amplitude $>1$. So, we search for divergences in the saddle-point equations for the propagator of the disorder-averaged generating functional to find conditions on $\boldsymbol{W}$ that lead to instability.

###### 4.8.3 Outlier eigenvalues of connectivity and relation to the propagator

While equations (S39)-(S42) hold for arbitrary mean connectivities, we in the following assume a homogeneous network with $\mu_{ij}=\mu$. In this case, the propagator drastically simplifies

$$\begin{aligned}\boldsymbol{K}^{-1} & = (1-P^{*}-T^{*})^{-1}\left(\mathbf{1}+\frac{\mu+\tilde{T}^{*}}{1-P^{*}-T^{*}-N(\mu+\tilde{T}^{*})}\{\mathbf{1}\}\right) & \text{(S43)}\\ & = (1-P^{*}-T^{*})^{-1}\left(\mathbf{1}+\mathcal{O}(1/N)\{\mathbf{1}\}\right). & \text{(S44)}\end{aligned}$$

From the saddle point equations, we then obtain

$$\begin{aligned}P^{*} & = B\frac{\sigma^{2}}{N}\sum_{i}\left[\boldsymbol{K}^{-1}\right]_{ii}=\frac{\left(\tau_{\mathrm{rec}}-2\tau_{\mathrm{chn}}\right)\sigma^{2}}{1-P^{*}-T^{*}}+\mathcal{O}(1/N),\\ \tilde{T}^{*} & = \tau_{\mathrm{chn}}\frac{\sigma^{2}}{N}\left\langle \tilde{\boldsymbol{X}}^{T}\boldsymbol{X}\right\rangle_{P^{*},Q^{*},T^{*}}=\frac{\tau_{\mathrm{chn}}\sigma^{2}}{1-P^{*}-T^{*}}+\mathcal{O}(1/N),\\ T^{*} & = \tau_{\mathrm{chn}}\frac{\sigma^{2}}{N}\sum_{i,j}\left[\boldsymbol{K}^{-1}\right]_{ij}=\frac{\tau_{\mathrm{chn}}\sigma^{2}}{1-P^{*}-T^{*}-N(\mu+\tilde{T}^{*})}=\mathcal{O}(1/N),\end{aligned}$$

so we can neglect $T^*$ and obtain

$$P^* = \frac{1}{2} \pm \sqrt{\frac{1}{4} - B\sigma^2}\,, \tag{S45}$$

$$\tilde{T}^* = \frac{\tau_{\mathrm{chn}}\sigma^2}{1 - P^*}\,. \tag{S46}$$

Only the solution of (S45) with the minus sign is physically plausible as $P^* = 0$ for $B = 0$.

In order to extract information about eigenvalues of the effective connectivity $\boldsymbol{W}$, we next analyze the divergence of $\boldsymbol{K}^{-1}$: Both $\mu$ and $\sigma$ depend linearly on the magnitude of the weights. Let's denote this scale with $\alpha$ and rescale $\mu \to \alpha\mu$ and $\sigma^2 \to \alpha^2\sigma^2$. The task is to find the scale of weights $\alpha$ that, for given parameters $\tau, \mu, \sigma$, leads to a divergence. In order to find that scale $\alpha$, we need to solve the divergence criterion in (S43)

$$0 = 1 - P^*(\alpha) - N\mu\alpha + \frac{N\tau_{\mathrm{chn}}\sigma^2\alpha^2}{1 - P^*(\alpha)} \tag{S47}$$

for $\alpha$. Here we neglected $T^*$ as this is $\mathcal{O}(1/N)$. The outlier eigenvalues $\lambda_o$ of $\boldsymbol{W}$ are then obtained by knowing that all eigenvalues $\lambda$ also scale with $\alpha$, so $\lambda \to \alpha\lambda$, and the divergence takes place if one eigenvalue is at least one. So we investigate the limit case $\alpha\lambda = 1$. Therefore, the outlier eigenvalues are given by $\lambda_o = \alpha^{-1}$. The general solution of (S47) for homogeneous networks is

$$\begin{aligned} \alpha &= \frac{(B - N\tau_{\mathrm{chn}})\,N\mu}{2\left((B + N\tau_{\mathrm{chn}})^2\,\sigma^2 + BN^2\mu^2\right)} \\ &\pm \frac{\sqrt{(-B + N\tau_{\mathrm{chn}})^2\,N^2\mu^2 + 4\left((B + N\tau_{\mathrm{chn}})^2\,\sigma^2 + BN^2\mu^2\right)N\tau_{\mathrm{chn}}}}{2\left((B + N\tau_{\mathrm{chn}})^2\,\sigma^2 + BN^2\mu^2\right)} \end{aligned}$$

which reduces to $\lambda_o = \alpha^{-1} = N\mu + \mathcal{O}(\tau_{\mathrm{rec}}/\sqrt{N})$ for $\tau_{\mathrm{chn}} = 0$. This special case shows that reciprocal motifs only have a minor effect on the outlier eigenvalues of $\boldsymbol{W}$, such that the dominant contribution is $\lambda_o = N\mu$, which is known from purely random inhibitory networks. In summary, outlier eigenvalues of $\boldsymbol{W}$ are created by non-zero mean connectivity as well as chain motifs.

To avoid instabilities from outlier eigenvalues of $\boldsymbol{W}$, the mean connectivity $\mu$ must be either negative, or positive but smaller than $1/N$ (see discussion below on excitatory and inhibitory networks). The above equations show that not only $\mu$ appears in combination with $N$, but the same is true for $\tau_{\mathrm{chn}}$. Therefore, if outliers are not close to the instability line and therefore leading to divergences, then their effect is small and can be neglected.

**Outlier eigenvalues of $\boldsymbol{W}$ and dimensionality** If outlier eigenvalues of $\boldsymbol{W}$ are close to the instability line, then their corresponding mode dominates the dynamics. At the critical point, the dynamics is effectively one dimensional [12]. This illustrates that the outlier eigenvalues of $\boldsymbol{W}$ shape the dimensionality in networks, where they are close to the instability line. In networks where the bulk of eigenvalues is closer to the instability line - the scenario we focus on here - the dimensionality is shaped by the spectral radius.

**Excitation-dominated networks** For excitatory networks, we have $\mu_{ij} > 0$, so the eigenvalue due to the mean connectivity could yield a divergence, especially in large networks when we use the above mentioned scaling of weights as $1/\sqrt{(N)}$. Therefore, for stable single-population excitatory networks, one would rather employ a scaling that yields $\mu_{ij} \sim 1/N$. The outlier eigenvalues of $\boldsymbol{W}$ are furthermore affected by $P^*, T^*, \tilde{T}^*$, i.e. reciprocal and chain motifs (cf. (S43)). Therefore, in excitation-dominated networks, only chain and reciprocal motifs affect the dimensionality, but not convergent and divergent motifs, in line with findings in [28].

**Inhibition-dominated networks** In inhibitory networks we can apply the standard standard weight scaling $W_{ij} \sim 1/\sqrt(N)$, so we have $\mu \sim \mathcal{O}(1/\sqrt{N}) < 0$ and thus no divergences due to the mean connectivity. In this case, we can ignore effects from the mean connectivity. Let us for the remainder also assume that there is no instability due to the chain outlier. Then the $\mathcal{O}(1/N)$ correction in (S43) is small and we can approximate $\boldsymbol{K}^{-1} = \frac{1}{1-P^*}\mathbf{1}$.

##### 4.8.4 Spectral radius and correlation functions

For the inhibitory network setting discussed above, we obtain from the saddle-point equations (S39) for the correlation function

$$Q^* = \frac{A\sigma^2}{(1-P^*)^2 - A\sigma^2} D. \tag{S48}$$

If the spectral radius $R$ of effective connectivity approaches one, then one eigendirection of the dynamics becomes unstable, and as a consequence correlations diverge. Since $Q^*$ measures correlations (cf. (S39)), the aim is to find a divergence of $Q^*$. From the equation above follows the condition

$$(1-P^*)^2 - A\sigma^2 = 0, \tag{S49}$$

which has to be rewritten in the following general form

$$R(\sigma^2, \tau_{\text{con}}, \tau_{\text{div}}, \tau_{\text{chn}}, \tau_{\text{rec}}) = 1 \tag{S50}$$

to infer the functional dependence of the spectral radius on the various second order statistics. Eigenvalues of $\boldsymbol{W}$ scale with the strength of connections $W_{ij}$ and therefore does the standard deviation $\sigma$. Hence, $R$ needs to be proportional to $\sigma$. Starting from (S49), we get

$$P^* + \sqrt{A}\sigma = 1 \tag{S51}$$

which, due to the fact that $P$ contains a square root, is not yet of the desired form (S50). Inserting (S45), squaring and reordering the resulting equation yields the desired form

$$\frac{A+B}{\sqrt{A}}\sigma = 1\,. \tag{S52}$$

The spectral radius can then be read off as

$$R = \frac{A+B}{\sqrt{A}}\sigma = \frac{1-\tau_{\text{div}}-\tau_{\text{con}}+\tau_{\text{rec}}-2\tau_{\text{chn}}}{\sqrt{1-\tau_{\text{div}}-\tau_{\text{con}}}}\sigma. \tag{S53}$$

The spectral radius is therefore affected by all motifs.

#### 4.9 Generalization to networks with multiple populations

In the following, we consider networks with multiple $(M)$ populations where the connection statistics to neurons only depends on the presynaptic populations. The latter constraint significantly reduces the parameter space. This is possible for all second order statistics except for the reciprocal motifs, as for the latter the postsynaptic neuron of the first connection is the presynaptic neuron of the second connection. For such connection statistics, we obtain

$$\begin{aligned}
\Delta_{ijkl} &= \langle\langle W_{ik}W_{jl}\rangle\rangle\, N \\
&= \sigma_k^2\delta_{ij}\delta_{kl} + \kappa_{\text{div},k}(1-\delta_{ij})\delta_{kl} + \kappa_{\text{con},kl}(1-\delta_{kl})\delta_{ij} \\
&+ \kappa_{\text{chn},lk}\delta_{il}(1-\delta_{jk}) + \kappa_{\text{chn},kl}\delta_{jk}(1-\delta_{il}) + \kappa_{\text{rec},kl}\delta_{il}\delta_{jk} \\
&= \delta_{ij}\delta_{kl}\left(\sigma_k^2 - \kappa_{\text{div},k} - \kappa_{\text{con},kl}\right) \\
&+ \delta_{il}\delta_{jk}\left(\kappa_{\text{rec},kl} - \kappa_{\text{chn},kl} - \kappa_{\text{chn},lk}\right) \\
&+ \text{terms containing a single}\,\delta\,,
\end{aligned}$$

where we can neglect the terms containing a single Kronecker delta (see discussion of (S38) above). Inserted in the action, we obtain a term

$$\begin{aligned}\sum_{i,j,k,l}\Delta_{ijkl}\tilde{X}_i\tilde{X}_jX_kX_l &= \sum_{i,k}\left(\sigma_k^2-\kappa_{\mathrm{div},k}-\kappa_{\mathrm{conv},kk}\right)\tilde{X}_i\tilde{X}_iX_kX_k\\ &+ \sum_{k,l}\left(\kappa_{\mathrm{rec},kl}-\kappa_{\mathrm{chain},kl}-\kappa_{\mathrm{chain},lk}\right)\tilde{X}_kX_k\tilde{X}_lX_l\\ &= \sum_i\tilde{X}_i\tilde{X}_i\sum_\alpha\underbrace{\left(\sigma_\alpha^2-\kappa_{\mathrm{div},\alpha}-\kappa_{\mathrm{conv},\alpha\alpha}\right)}_{A_\alpha}\sum_{k\in\alpha}X_kX_k\\ &+ \sum_{\alpha,\beta}\underbrace{\left(\kappa_{\mathrm{rec},\alpha\beta}-\kappa_{\mathrm{chain},\alpha\beta}-\kappa_{\mathrm{chain},\beta\alpha}\right)}_{B_{\alpha\beta}=B_{\beta\alpha}}\sum_{k\in\alpha}\tilde{X}_kX_k\sum_{l\in\beta}\tilde{X}_lX_l\end{aligned}$$

where the sum over $\alpha$ and $\beta$ run over all $M$ populations. Now we insert auxiliary fields

$$\begin{aligned}Q_\alpha &= \frac{1}{N_\alpha}\sum_{k\in\alpha}X_kX_k\,,\\ P_\alpha &= \frac{1}{N_\alpha}\sum_{k\in\alpha}\tilde{X}_kX_k\,,\end{aligned}$$

and write

$$\begin{aligned}1 &\sim \int DQ_\alpha\int D\tilde{Q}_\alpha\,\exp\left(-N_\alpha Q_\alpha\tilde{Q}_\alpha+\tilde{Q}_\alpha\textstyle\sum_{k\in\alpha}X_kX_k\right)\,,\\ 1 &\sim \int DP_\alpha\int D\tilde{P}_\alpha\,\exp\left(-N_\alpha P_\alpha\tilde{P}_\alpha+\tilde{P}_\alpha\textstyle\sum_{k\in\alpha}\tilde{X}_kX_k\right)\,,\end{aligned}$$

so that we get a disorder-averaged generating function (for simplicity we set $\boldsymbol{\mu}=0$ which is a good approximation as long as $\boldsymbol{\mu}$ is not tuned to drive the system close to instability, see above)

$$\begin{aligned}\left\langle\tilde{Z}(\mathbf{J})\right\rangle_W &\sim \int\boldsymbol{DP}\int\boldsymbol{DQ}\exp\left(S(\boldsymbol{P},\boldsymbol{Q})\right)\\ S(\boldsymbol{P},\boldsymbol{Q}) &= -\sum_\alpha\left(N_\alpha Q_\alpha\tilde{Q}_\alpha+N_\alpha P_\alpha\tilde{P}_\alpha\right)+\frac{1}{2N}\sum_{\alpha,\beta}B_{\alpha\beta}N_\alpha P_\alpha N_\beta P_\beta+\ln\Omega\left(\boldsymbol{P},\boldsymbol{Q}\right)\\ \Omega\left(\boldsymbol{P},\boldsymbol{Q}\right) &= \int D\mathbf{X}\int D\tilde{\boldsymbol{X}}\exp\left(-\tilde{\boldsymbol{X}}^T\left(\mathbf{1}-\tilde{\boldsymbol{P}}\right)\boldsymbol{X}+\frac{1}{2}\left(D+\frac{\sum_\alpha N_\alpha Q_\alpha A_\alpha}{N}\right)\tilde{\mathbf{X}}^T\tilde{\mathbf{X}}+\boldsymbol{X}^T\tilde{\boldsymbol{Q}}\boldsymbol{X}+\mathbf{J}\mathbf{X}\right)\,.\end{aligned}$$

As for fully random networks, we evaluate the exponent of the disorder-averaged generating function in saddle-point approximation, which yields

$$\tilde{Q}_\alpha^*\sim\left\langle\tilde{X}_i\tilde{X}_i\right\rangle=0$$

and, in addition,

$$Q_\alpha^* = \frac{1}{N_\alpha}\left\langle \sum_{k\in\alpha} X_k X_k \right\rangle_{P^*,Q^*} = \frac{1}{N_\alpha}\sum_{k\in\alpha}\left[\left(\mathbf{1}-\tilde{\boldsymbol{P}}^*\right)^{-1}\left(D+\frac{\sum_\gamma N_\gamma Q_\gamma A_\gamma}{N}\right)\left(\mathbf{1}-\tilde{\boldsymbol{P}}^*\right)^{-T}\right]_{kk} = \frac{D+\frac{\sum_\gamma N_\gamma Q_\gamma A_\gamma}{N}}{(1-\tilde{P}_\alpha^*)^2} \tag{S54}$$

$$P_\alpha^* = \frac{1}{N_\alpha}\left\langle \sum_{k\in\alpha} \tilde{X}_k X_k \right\rangle_{P^*,Q^*} = \frac{1}{N_\alpha}\sum_{k\in\alpha}\left[\left(\mathbf{1}-\tilde{\boldsymbol{P}}^*\right)^{-1}\right]_{kk} = \frac{1}{1-\tilde{P}_\alpha^*} \tag{S55}$$

$$\tilde{P}_\alpha^* = \frac{1}{N}\sum_\beta B_{\alpha\beta} N_\beta P_\beta^* \tag{S56}$$

Solving (S54) yields

$$Q_\alpha^* = \sum_\beta M_{\alpha,\beta} P_\alpha^{*2} D,$$

with $M_{\alpha,\beta} = \delta_{\alpha,\beta} - P_\alpha^{*2}\frac{N_\beta}{N}A_\beta$. We obtain a divergence if $\det(M) = 0$. Since $M$ is the identity matrix shifted by an outer product, we can use the matrix determinant lemma to calculate the determinant of $M$:

$$\det M = 1 - \sum_\gamma \frac{N_\gamma}{N} A_\gamma P_\gamma^{*2} = 0 \tag{S57}$$

which shows a divergence, similar as in the case for the single-population network above. In fact, the divergence expression reduces to that of the single-population network for homogeneous $\tilde{P}_\alpha^* = \tilde{P}^*$, which follows from homogeneous $B_{\alpha\beta} = B$ for all $\alpha, \beta$. If the latter equality does not hold, then finding the spectral radius amounts to solving a set of coupled nonlinear equations, which gives very convoluted expressions. We therefore here approximate $B_{\alpha\beta} \approx \frac{1}{M}\sum_\alpha B_{\alpha\beta} = B_\beta$ and obtain from (S55) and (S56)

$$P^* = \frac{1}{1-\sum_\beta B_\beta \frac{N_\beta}{N} P^*}. \tag{S58}$$

Note that an alternative approximation would be a weighted average $B_{\alpha\beta} \approx \sum_{\alpha,\beta}\frac{N_\alpha}{N}B_{\alpha\beta} = B_\beta$ yielding qualitatively similar results. Equations (S57) and (S58) then have the same form as in the single-population network and can be solved analogously. Redefining

$$\begin{aligned}
\sigma^2 &= \sum_\alpha \frac{N_\alpha}{N}\sigma_\alpha^2, \\
\tau_{\text{div}} &= \frac{\kappa_{\text{div}}}{\sigma^2} = \frac{\sum_\alpha \frac{N_\alpha}{N}\kappa_{\text{div},\alpha}}{\sum_\alpha \frac{N_\alpha}{N}\sigma_\alpha^2}, \\
\tau_{\text{con}} &= \frac{\kappa_{\text{con}}}{\sigma^2} = \frac{\sum_\alpha \frac{N_\alpha}{N}\kappa_{\text{con},\alpha\alpha}}{\sum_\alpha \frac{N_\alpha}{N}\sigma_\alpha^2}, \\
\tau_{\text{rec}} &= \frac{\kappa_{\text{rec}}}{\sigma^2} = \frac{\frac{1}{M}\sum_{\alpha,\beta}\frac{N_\beta}{N}\kappa_{\text{rec},\alpha\beta}}{\sum_\alpha \frac{N_\alpha}{N}\sigma_\alpha^2}, \\
\tau_{\text{chn}} &= \frac{\kappa_{\text{chn}}}{\sigma^2} = \frac{\frac{1}{2M}\sum_{\alpha,\beta}\frac{N_\alpha+N_\beta}{N}\kappa_{\text{chn},\alpha\beta}}{\sum_\alpha \frac{N_\alpha}{N}\sigma_\alpha^2}
\end{aligned}$$

as combinations of motif statistics of multiple populations, we obtain the same expression for the spectral radius

$$R = \frac{1-\tau_{\text{div}}-\tau_{\text{con}}+\tau_{\text{rec}}-2\tau_{\text{chn}}}{\sqrt{1-\tau_{\text{div}}-\tau_{\text{con}}}}\sigma. \tag{S59}$$

#### 4.10 From covariances of connections to motif abundances

Due to the limited quantity of data on the effective strengths of synapses in the synaptic physiology datasets, we quantified only binarized motif statistics the data. Here we demonstrate how to translate between cumulants $\kappa$ and binarized motif statistics for an example network of one excitatory and one inhibitory population. In the framework above all excitatory cells have the same strengths $J_E = J$ and all inhibitory synapses have the same strengths $J_I = -gJ$. In excitatory-inhibitory networks, the typically larger number of excitatory neurons is often compensated by a larger strength of inhibitory synapses (see [29, 20]). In going from covariances of connections to correlation coefficients $\tau$, the overall synaptic strength $J$ drops out, but the relative strength $-g$ remains. For excitatory connection probabilities $p_E$, inhibitory connection probabilities $p_I$ and a ratio $\gamma = N_I/N_E$ between inhibitory and excitatory population sizes, we obtain

$$\begin{aligned}
\tau_{\text{div}} &= \frac{\hat{\sigma}_E^2 \tau_{\text{div},E} + \gamma g^2 \hat{\sigma}_I^2 \tau_{\text{div},I}}{\hat{\sigma}_E^2 + \gamma g^2 \hat{\sigma}_I^2}, \\
\tau_{\text{con}} &= \frac{\hat{\sigma}_E^2 \tau_{\text{con},EE} + \gamma g^2 \hat{\sigma}_I^2 \tau_{\text{con},II}}{\hat{\sigma}_E^2 + \gamma g^2 \hat{\sigma}_I^2}, \\
\tau_{\text{rec}} &= \frac{1}{2} \frac{\hat{\sigma}_E^2 \tau_{\text{rec},EE} - (1+\gamma) g \sqrt{\hat{\sigma}_E^2 \hat{\sigma}_I^2}\, \tau_{\text{rec},EI} + \gamma g^2 \hat{\sigma}_I^2 \tau_{\text{rec},II}}{\hat{\sigma}_E^2 + \gamma g^2 \hat{\sigma}_I^2}, \\
\tau_{\text{chn}} &= \frac{1}{2} \frac{\hat{\sigma}_E^2 \tau_{\text{chn},EE} - \frac{1+\gamma}{2} g \sqrt{\hat{\sigma}_E^2 \hat{\sigma}_I^2}\, (\tau_{\text{chn},EI} + \tau_{\text{chn},IE}) + \gamma g^2 \hat{\sigma}_I^2 \tau_{\text{chn},II}}{\hat{\sigma}_E^2 + \gamma g^2 \hat{\sigma}_I^2},
\end{aligned}$$

with $\hat{\sigma}_E^2 = p_E(1-p_E)$ and $\hat{\sigma}_I^2 = p_I(1-p_I)$.

#### 4.11 Constraints on motif abundances and algorithms for network construction

Given that the spectral radius is affected by all motifs, we now ask the question, which abundances of motifs are possible. We already saw one restriction above, namely that in homogeneous networks $\tau_{\text{chn}}$ must be either on the order $\mathcal{O}(1/N)$ or negative in order to yield stable dynamics (see discussion on outlier eigenvalues above). However, there are more restrictions on the motif abundances coming from the fact that the covariance matrix of connections weights must be positive definite: no negative eigenvalues are allowed. Reciprocal motifs always yield positive definite covariance matrices, therefore all values $\tau_{\text{rec}} \in [-1, 1]$ are possible.

The situation is different for divergent and convergent motifs: while any positive value $\tau \in [0, 1]$ is possible, only negative values $> -1/N$ are allowed. An intuitive explanation for this is that for divergent and convergent motifs any connection is not only part of one possible motif - as it is the case for reciprocal motifs - but of $N$ motifs (here we also count motifs with self-connections). In order to have negative values for $\tau_{\text{div}}$ or $\tau_{\text{con}}$, pairs of connections must be negatively correlated. For Gaussian networks, this can be achieved by drawing a common Gaussian distributed number $\alpha_i$ and setting $W_{ij} \leftarrow W_{ij} + \alpha_i$ and $W_{ik} \leftarrow W_{ik} - \alpha_i$ for one $k \neq i$. In order for the connection $W_{ij}$ to be negatively correlated to all $k \neq i$, one would need to draw a new number $\alpha_i$ for each neuron $k$, therefore in total $N$ numbers. These $N$ numbers of course also contribute to the variance of connection $W_{ij}$. Therefore each number $\alpha_i$ can only have a variance of order $1/N$ and therefore $\tau$ can only be of order $1/N$. One might think that the same situation applies for positive $\tau$. However, in this case it is possible to draw a single number $\alpha_i$ for all $k$. This also creates a higher-order motif, but also leads to the fact that all pairs of connections ending at neuron $i$ are positively correlated. Therefore, any value $\tau \in [0, 1]$ is possible for convergent motifs. The same rationale holds for divergent motifs.

We find again another situation for chain motifs: if one wants to include chains only, then the range of $\tau_{\text{chn}}$ is restricted to values $|\tau_{\text{chn}}| < 1/N$. For positive $\tau_{\text{chn}}$, this is in line with the stability issue of the outlier eigenvalue above. But for negative $\tau_{\text{chn}}$ we find the same argument as for convergent and divergent motifs above: each connection is part of $\mathcal{O}(N)$ chains. Negative correlations of sizable strength can therefore not be realized. However, there is another possibility. If one allows creation of divergent, convergent and reciprocal motifs while creating chains, one possibility is to draw one random number $\alpha_i$ for all connections that start or end at neuron $i$. This creates chain correlations, but also divergent, convergent and reciprocal ones. In this case, one can obtain a positive definite covariance matrix for larger values of $\tau_{\rm chn}$, but large positive values are additionally ruled out by the outlier instability. Therefore, only one possible case remains, where a single $\alpha_i$ is drawn for all connections starting and ending at neuron $i$, but $W_{ij} \leftarrow W_{ij} + \alpha_i$ and $W_{ji} \leftarrow W_{ji} - \alpha_i$ for all $j$ to create negative chain correlations.

The above considerations lead to the following algorithms for the creation of Gaussian networks with desired second-order statistics (see also [16]): We draw weights

$$W_{ij} = \mu + A\nu_{ij} + B\nu_{ji} + C\eta_i + D\eta_j$$

with independent and identically distributed variables $\nu_{ij} \overset{i.i.d.}{\sim} \mathcal{N}(0,1)$ and $\eta_i \overset{i.i.d.}{\sim} \mathcal{N}(0,1)$, so that coefficients $A, B, C, D$ encode covariances between weights

$$\begin{aligned}
\langle W_{ij}W_{kl}\rangle - \langle W_{ij}\rangle\langle W_{kl}\rangle &= \langle (A\nu_{ij} + B\nu_{ji} + C\eta_i + D\eta_j)\,(A\nu_{kl} + B\nu_{lk} + C\eta_k + D\eta_l)\rangle \\
&= \left(A^2 + B^2\right)\delta_{ik}\delta_{jl} + 2AB\delta_{il}\delta_{jk} \\
&+ C^2\delta_{ik} + D^2\delta_{jl} + CD\delta_{il} + CD\delta_{jk} \\
&= \underbrace{\left(A^2 + B^2 + C^2 + D^2\right)}_{=\sigma^2/N}\delta_{ik}\delta_{jl} + \underbrace{(2AB + 2CD)}_{=\sigma^2\tau_{\rm rec}/N}\delta_{il}\delta_{jk} \\
&+ \underbrace{C^2}_{=\sigma^2\tau_{\rm con}/N}\delta_{ik}(1-\delta_{jl}) + \underbrace{D^2}_{=\sigma^2\tau_{\rm div}/N}\delta_{jl}(1-\delta_{ik}) \\
&+ \underbrace{CD}_{=\sigma^2\tau_{\rm chn}/N}\left(\delta_{il}(1-\delta_{jk}) + \delta_{jk}(1-\delta_{il})\right)\,.
\end{aligned}$$

Solving the above relations yields equations for the coefficients $A, B, C, D$ in terms of the motif statistics:

$$\begin{aligned}
A &= \frac{\sigma}{2\sqrt{N}}\sqrt{1-\tau_{\rm con}-\tau_{\rm div}}\left(\sqrt{1+U}+\sqrt{1-U}\right)\,, \\
B &= \frac{\sigma}{2\sqrt{N}}\sqrt{1-\tau_{\rm con}-\tau_{\rm div}}\left(\sqrt{1+U}-\sqrt{1-U}\right)\,, \\
C &= \pm\frac{\sigma}{\sqrt{N}}\sqrt{\tau_{\rm con}}\,, \\
D &= \pm\frac{\sigma}{\sqrt{N}}\sqrt{\tau_{\rm div}}\,,
\end{aligned}$$

with

$$U = \frac{\tau_{\rm rec} - 2\tau_{\rm chn}}{1-\tau_{\rm con}-\tau_{\rm div}}\,.$$

We obtain the restriction $\tau_{\rm chn} = \pm\sqrt{\tau_{\rm con}\tau_{\rm div}}$, i.e. chain motifs can only be created along with convergent and divergent motifs, as previously reported in [17]. To avoid positive outlier eigenvalues of $\boldsymbol{W}$, in the numerical simulations presented in the main text, we mostly chose $C = \frac{\sigma}{\sqrt{N}}\sqrt{\tau_{\rm con}}$, and $D = -\frac{\sigma}{\sqrt{N}}\sqrt{\tau_{\rm div}}$ such that $\tau_{\rm chn} = -\sqrt{\tau_{\rm con}\tau_{\rm div}} < 0$. Since values for $U$ are restricted to the interval $[-1, 1]$ we furthermore chose $\tau_{\rm rec} = 2\tau_{\rm chn}$ and $\tau_{\rm con} = \tau_{\rm div}$ to increase the range of possible values $\tau_{\rm chn}$.

For excitatory-inhibitory networks, the coefficients receive population indices $\alpha, \beta, \gamma, \delta \in \{E, I\}$. For $i \in \alpha$, $j \in \beta$, $k \in \gamma$ and $l \in \delta$, we draw weights $W_{ij} = \mu + A_{\alpha\beta}\nu_{ij} + B_{\alpha\beta}\nu_{ji} + C_\beta\eta_i + D_\beta\eta_j$ and obtain the following coefficients

$$
\begin{aligned}
A_{\alpha\beta} &= \frac{\sigma_\beta}{2\sqrt{N}}\sqrt{1-\tau_{\mathrm{con},\beta\beta}-\tau_{\mathrm{div},\beta}}\left(\sqrt{1+U_{\alpha\beta}}+\sqrt{1-U_{\alpha\beta}}\right)\\
B_{\alpha\beta} &= \frac{\sigma_\beta}{2\sqrt{N}}\sqrt{1-\tau_{\mathrm{con},\beta\beta}-\tau_{\mathrm{div},\beta}}\left(\sqrt{1+U_{\alpha\beta}}-\sqrt{1-U_{\alpha\beta}}\right)\\
C_\alpha &= \pm\frac{\sigma_\alpha}{\sqrt{N}}\sqrt{\tau_{\mathrm{con},\alpha\alpha}}\,,\\
D_\alpha &= \pm\frac{\sigma_\alpha}{\sqrt{N}}\sqrt{\tau_{\mathrm{div},\alpha}}\,,
\end{aligned}
$$

with

$$
U_{\alpha\beta} = (-1+2\delta_{\alpha\beta})\,\frac{\tau_{\mathrm{rec},\alpha\beta}-\tau_{\mathrm{chn},\alpha\beta}-\tau_{\mathrm{chn},\beta\alpha}}{\sqrt{1-\tau_{\mathrm{con},\alpha\alpha}-\tau_{\mathrm{div},\alpha}}\sqrt{1-\tau_{\mathrm{con},\beta\beta}-\tau_{\mathrm{div},\beta}}}\,.
$$

Note that the overall minus sign in $U_{EI}$ arises from a relative minus sign between correlation coefficients $\tau_{\mathrm{rec},EI}$ of motif abundances (binarized statistics) and covariances $\kappa_{\mathrm{rec},EI}$ that include synaptic weights, i.e. $\kappa_{\mathrm{rec},EI} = -gJ^2\left(\sqrt{\hat{\sigma}_E^2\hat{\sigma}_I^2}\tau_{\mathrm{rec},EI}\right) = -\sigma_E\sigma_I\tau_{\mathrm{rec},EI}$. The same holds true for chain motifs between excitatory and inhibitory neurons, $\kappa_{\mathrm{chn},\alpha\beta} = -\sigma_E\sigma_I\tau_{\mathrm{rec},\alpha\beta}$ for $\alpha \neq \beta$. We obtain the restriction $\tau_{\mathrm{chn},\alpha\alpha} = -\sqrt{\tau_{\mathrm{con},\alpha\alpha}\tau_{\mathrm{div},\alpha}}$ and $\tau_{\mathrm{chn},\alpha\beta} = +\sqrt{\tau_{\mathrm{con},\alpha\alpha}\tau_{\mathrm{div},\beta}}$ for $\alpha \neq \beta$.
